# Spatially resolved profiling of protein conformation and interactions by biocompatible chemical cross-linking in living cells

**DOI:** 10.1101/2022.01.20.476705

**Authors:** Lili Zhao, Qun Zhao, Yuxin An, Hang Gao, Weijie Zhang, Zhou Gong, Xiaolong Liu, Baofeng Zhao, Zhen Liang, Chun Tang, Lihua Zhang, Yukui Zhang

## Abstract

The protein structures and interactions that maintain and regulate cellular processes in different subcellular organelles are heterogeneous and dynamic. However, it remains challenging to characterize the subcellular specificity and translocation of protein complexes in terms of conformation and interactions. Herein, we developed a spatially resolved protein complex profiling approach by biocompatible chemical cross-linking in living cells (SPACX) to monitor the dynamics of protein conformation, interactions and translocation. The advancement of fast capturing protein complexes in the physiological state, coupled with efficient enrichment of the cross-linked peptides, ensured deep-coverage analysis of the protein interactome in living cells. By ensemble structure refinement with cross-linking restraints, subcellular-specific conformation heterogeneity was identified for PTEN. PTEN displayed a broader range of dynamic conformation changes on the dual specificity domains in the nucleus than in the cytoplasm. Moreover, based on conformational differences, different interacting assemblies involving 25 cytoplasm-exclusively and 18 nucleus-exclusively PTEN-interacting proteins were found to account for diverse biological functions. Upon ubiquitin-proteasome system (UPS) stress, the assembly of PTEN and its interacting partners changed obviously during translocation. We newly identified 36 PTEN-interacting proteins, which were found to be highly enriched in functional pathways closely related to cell apoptosis. Inspiringly, the interactions among PTEN isoforms and their interacting proteins were accessible by the determination of sequence-unique cross-linking interfaces for direct interactions. All these results indicate the promise of SPACX to elucidate the functional heterogeneity of proteins in individual subcellular sociology.

## Introduction

Proteins assemble with interacting partners in a sophisticated manner by undergoing conformational changes for functional regulations. The cellular environment significantly affects the structure and interactions of protein complexes (1, 2). Unlike in vitro conditions, cellular environment has the typical features of crowding, sticking and confinement effects, which are indispensable for maintaining protein stability and aggregation (3). In addition, high-dimensional signal transduction network is formed by the dynamic assembly of protein complexes shuttling through different subcellular spaces to enable exclusive structures and interactions for specific functions (4). Thus, characterization of the spatially dynamic conformation and interaction of protein complexes in living cells is highly desirable.

Many technologies have been developed to study protein structures and interactions in a cellular context. Cryo-electron tomography (cryo-ET) allows the generation of the architectural structure of giant protein machines such as the nuclear pore complex (5, 6), which are difficult to purify intact in vitro. In-cell nuclear magnetic resonance spectroscopy (in-cell NMR) improved the dynamic resolution of proteins in the native crowded cellular environment (7, 8). However, these approaches could not satisfy the requirement for proteome-wide protein characterization. Recently, proximity labeling technology was proposed to capture the spatially adjacent protein interactions in living cells. A BioID-based map of a human cell on the basis of 192 subcellular markers was built, and 35,902 interactions were established with 4,424 unique high-confidence proximity interactors (9). Although proximity labeling technology (10–12) is superior to affinity purification-mass spectrometry (AP-MS) (13, 14) in spatial resolution and weak interaction capture, the shuttle-dynamic changes in different organelles are still challenging to obtain, and cell engineering is needed for both methods.

To advance protein complex analysis with direct interaction interfaces, as well as protein structural constraints, chemical cross-linking mass spectrometry (CXMS) technology has become an indispensable supplement to the above methods (15, 16). This technology can freeze protein conformation and interactions via the formation of covalent bonds and allow global identification of the intra- and inter-cross-links of protein complexes with high sensitivity. Until now, CXMS has been successfully employed to analyze protein complexes from purified proteins (17, 18) and cell lysates (19–21). With further consideration of the effect of the cellular environment on protein structure and interactions, in vivo CXMS strategy was proposed. Huang group integrated a multifunctional MS-cleavable cross-linker with sample preparation strategies, enabling the construction of the largest in vivo cross-linking human PPI network to date (22). Bruce group used in vivo Protein Interaction Reporter (PIR) cross-linking to capture the protein interactions of membrane proteins to gain insight into bacterial antibiotic resistance mechanisms (23). However, the biocompatibility of chemical cross-linking in terms of cell viability and protein stability has not been carefully assessed to verify minimal disturbance to the physiological state in living cells.

Moreover, the heterogeneity of protein complexes within various subcellular organelles is largely homogenized in overall cellular results. Responsively, to study the protein interactome in specific subcellular organelles, Heck group used DSSO to crosslink separated human cell nuclei, revealing an overview of chromatin-associated PPIs (24). Borchers group utilized the enrichable cross-linker CBDPS to cross-link isolated yeast mitochondria, allowing high-throughput detection of PPIs involving a sixth of the mitochondrial proteome (25). Nevertheless, these approaches were still unable to characterize the dynamic assembly of protein translocation between different subcellular spaces, which was of interest to discover the biological functions of protein complexes in highly dynamic transitions.

In this work, we used a compact enrichable cross-linker to develop a spatially resolved protein complex profiling approach by biocompatible and deep-coverage chemical cross-linking in living cells (SPACX, Fig. 1). As a proof-of-concept, the spatially specific assemblies of PTEN within the cytoplasm and nucleus were confidently identified. Furthermore, the dynamic assembly of PTEN translocation from the cytoplasm to the nucleus was successfully obtained, allowing an overall view to illuminate the functions in which PTEN participates. Therefore, all these results demonstrated the utility of our method for achieving dynamic protein assemblies in living cells with good spatial resolution.

**Fig. 1.**
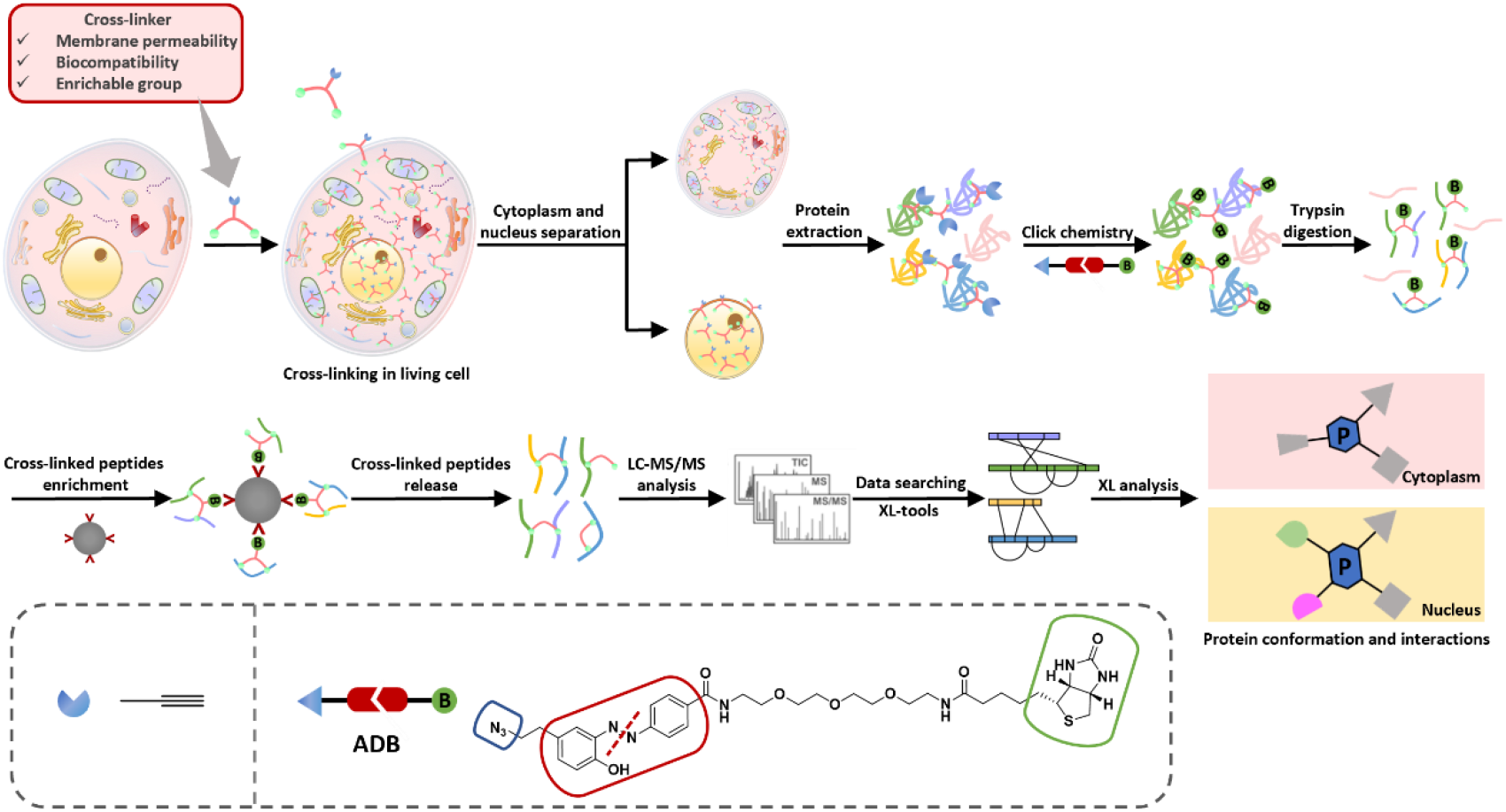
Schematic illustration of SPACX. Living cells are cross-linked in vivo and then the cytoplasm and nucleus are separated. The cross-linked proteins are extracted and reacted with ADB by bioorthogonal alkyne-azido click chemistry for biotin labeling. The biotin-labeled cross-linked proteins are subsequently digested into a peptide mixture, and the cross-linked peptides are affinity purified with streptavidin and released by a reduction reaction. Finally, the cross-linked peptides are subjected to MS identification for spatially specific cross-links analysis.

## Results

### Biocompatibility of cross-linking in living cells at minutes scale

To profile protein complexes in living cells by chemical cross-linking with high biocompatibility, the cross-linking reagent itself first needs to possess fast membrane permeability and high reactivity with two reactive amino acids located in protein complexes within the distance constraint of the cross-linker arm length. Then, good water solubility of the cross-linker is desired to minimize damage to the cell activity caused by the addition of an excess of organic reagents and to increase the amount of cross-linker used in the reaction. In addition, an enrichable moiety must be introduced to the cross-linker to improve the identification sensitivity of low-abundance cross-linked peptides by eliminating the interference of regular peptides and matrix. Notably, the cellular physiological state is essential to maintain; thus, the cross-linking reaction should ensure minimal interference with cell viability and protein stability without compromising the cross-linking coverage.

Given the high reactivity and reaction specificity of N-hydroxysuccinimidyl (NHS) ester for lysine residues, which are highly abundant on the protein surface, the cross-linker bis(succinimidyl)propargyl with nitro compound (BSPNO) was synthesized (*SI* Note S1 and Fig. S1-S9). BSPNO consists of two NHS reactive groups and one flexible enrichment group alkyne, with a maximum Cα–Cα distance restraint of 27 Å (*SI* Note S2). The basic properties, water solubility and size of BSPNO were evaluated by comparing cLogP and surface area values with those of currently popular cross-linkers (*SI* Note S3). As shown in Fig. 2A, the water solubility of BSPNO is better than that of the enrichable cross-linkers PIR (26) and Azide-A-DSBSO (27) and comparable to that of the commonly used highly water soluble cross-linker BS3 with impermeable sulfonic acid groups. Simultaneously, BSPNO is only slightly larger than the linear chain cross-linkers DSS, BS3 and DSSO (28), which lack enrichable capability, and much smaller than other enrichable cross-linkers. Hence, the suitable water solubility and size give BSPNO good solubility and membrane permeability in physiological buffer (1X PBS, pH 7.4; 1% DMSO added).

**Fig. 2.**
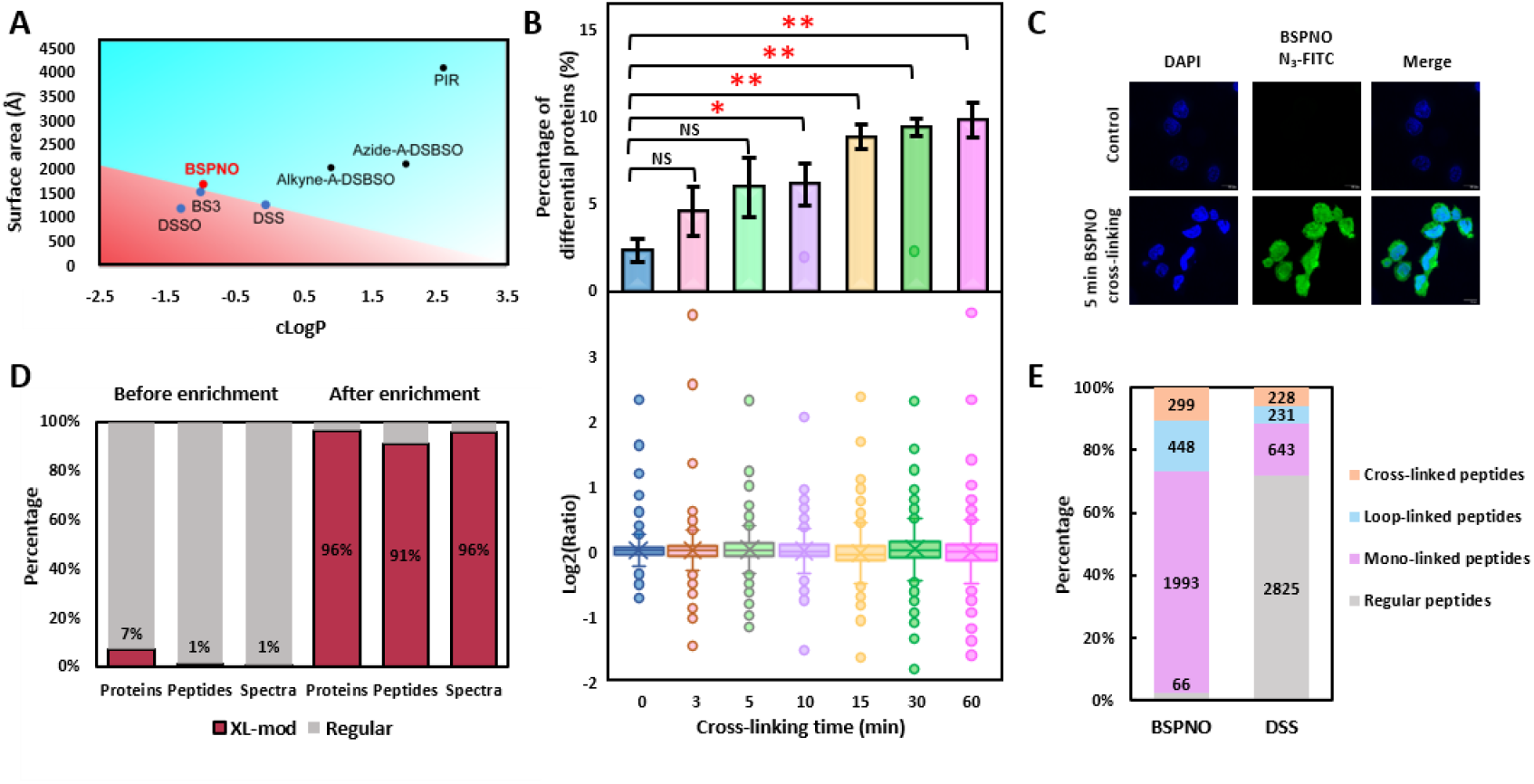
Assessment and construction of SPACX. (*A*) Surface area and cLogP for BSPNO and a series of widely used cross-linking reagents. (*B*) Upper: Histogram of the percentages of differentially expressed proteins for the BSPNO cross-linked cells at different cross-linking time. Bottom: Box plot of the integral Log2 ratio distribution for the corresponding cross-linking time. The p values were calculated based on the two-sided T-test. *p < 0.05, **p < 0.01, and ns indicates not significant. p (3 min) = 0.20; p (5 min) = 0.11; p (10 min) = 0.03; p (15 min) = 0.004; p (30 min) = 0.006; p (60 min) = 0.01. (*C*) Fluorescence imaging of BSPNO distribution in 5 min cross-linked cells. (*D*) Enrichment selectivity of cross-links by comparative analysis of the identified cross-links proportions among proteins, peptides and spectra using original and enriched cross-linked cell samples. (*E*) Percentage distributions of BSPNO and DSS cross-linked peptides.

Based on these properties, the influence of cross-linking time on cell viability and protein stability was investigated (*SI* Fig. S10). After 0 min, 3 min, 5 min, 10 min, 15 min, 30 min and 60 min of cross-linking with BSPNO, CCK-8 measurements indicated no obvious disturbance of cell activity (*SI* Note S4 and Fig. S11). The size of the cross-linked cells tended to shrink, as monitored by microscopy (*SI* Note S5 and Fig. S12). Most importantly, we evaluated the proteome stability of the cross-linked cells with different cross-linking time using a quantitative proteomics approach (*SI* Note S6). As shown in Fig. 2B, within 5 min of cross-linking, no obvious protein expression difference was shown. When the cross-linking time was extended, the percentage of differential proteins significantly increased, especially at the commonly adopted chemical cross-linking time of 60 min (27, 29). Furthermore, the influence of the above cross-linking time on protein structure stability was analyzed using bovine serum albumin (BSA), which has a relatively rigid structure, as a model. Interestingly, the number of identified cross-linked sites with Cα–Cα distances exceeding the maximum distance restraint of 27 Å obviously decreased with increasing cross-linking time: a 15% reduction in the occupied proportion was observed by comparing 60 min to 5 min of cross-linking, which implies that the structure of BSA might be locally unfolded due to covalent labeling perturbation (30, 31) (*SI* Note S7 and Fig. S13, S14).

In summary, as the cross-linking time increased, there was no significant disturbance in cell activity and morphology. However, protein expression in living cells was gradually affected, and protein structure was constricted, resulting in adverse induction of cellular homeostasis. Therefore, 5 min was chosen as the optimal BSPNO living cell cross-linking time in view of the minimal perturbation to the cellular physiological state. Moreover, the cross-linking effect was demonstrated by fluorescence imaging. BSPNO could be effectively transported inside the cells and cross-link intracellular proteins within 5 min (Fig. 2C, *SI* Note S8), which was crucial to fast capture the dynamic changes in intracellular protein complex assemblies with temporal resolution at the minute scale.

### Coverage of cross-linking in living cells

The essential prerequisite for proteome-wide CXMS study was to enhance the detection coverage of cross-linked products by enrichment. A cleavable azide-diazo-biotin (ADB) ligand was grafted to the cross-linked peptides by click chemistry of alkynyl derivatization to allow enrichment with streptavidin beads (*SI* Fig. S15 and Note S9). The sandwich method could compact the structure of the cross-linker for efficient membrane permeability and enable highly selective enrichment of the cross-linked peptides.

By enrichment, the identified cross-linker-modified proteins, peptides and spectra were all significantly improved (Fig. 2D). Compared with the commonly used nonenrichable cross-linker DSS, BSPNO increased the proportion of cross-linker-modified peptides identified from *E. coli* lysate, contributed by reducing the regular peptides from 72% to 2% (Fig. 2E, *SI* Note S10 and Fig. S16). In addition, more b/y fragment ions were matched with higher sequence continuity in the MS2 spectra identified with BSPNO than DSS (*SI* Fig. S17), and the spectra number matched for each of the cross-linked peptides also increased (*SI* Fig. S18), ensuring good accuracy for cross-links identification. Furthermore, similar abundance distribution range was observed for the cross-linked proteome and human MS proteome across seven orders of magnitude (*SI* Fig. S19). All these results indicated that in-depth analysis of the cross-linked peptides could be achieved by efficient enrichment coupled with biocompatible cross-linking in vivo, which was necessary prerequisite for conformational and interactional dynamics characterization.

### Proteome-wide validation of the cross-linking PPIs and structural mapping in living cells

We globally acquired the protein interactions and structural constraints in human BEL7402 cells, contributed by the high biocompatibility and enrichment efficiency of the cross-links. In total, 18,886 interprotein cross-linked sites (Dataset S1) were identified, mapping 3,366 interprotein interactions involved in 1,595 proteins. The constructed interprotein interaction network is presented in Fig. 3A (Dataset S2). These PPIs were associated with many functional proteins, such as histones, heat shock proteins (HSPs), transcription factors (TFs), ribosome proteins, zinc finger proteins, and calcium-related proteins, indicating the good ability of our cross-linking in living cells for interaction analysis of functional proteins. Further cellular component analysis showed that the PPIs were widely distributed throughout the cell, including the nucleus, cytosol, membrane, endoplasmic reticulum, and mitochondria. They extensively participate in various biological processes, including RNA transcription, signal transduction and cell adhesion processes, with molecular functions of nucleic acid binding, ATP binding, metal ion binding, and protein binding (*SI* Fig. S20). Importantly, living cell cross-linking showed great superiority in detecting histone protein interactions, which were easily destroyed in the process of cell lysis; heat shock protein interactions, which were typically weak and dynamic molecular chaperones; and relatively low-abundance transcription factor protein interactions (*SI* Fig. S21, S22, Note S11, Dataset S3). These accomplishments were attributed to the higher protein concentration provided by the unique crowding and confinement of the intracellular environment, which obviously contrasts with the dispersal of proteins in cell lysate. In addition, the abundance correlation of the pairs of interacting proteins was investigated, and the results showed that the interactions between proteins with different abundances across seven orders of magnitude could be efficiently captured and identified (*SI* Fig. S23). Then, this PPI dataset was matched with the existing PPI databases, and 61% complementarity (2,065 PPIs) was observed (*SI* Fig. S24, Dataset S2), which was most likely attributed to the capture bias and distinct filter threshold of these PPI profiling methods, as well as cell type and cell state heterogeneity. Next, we investigated the accuracy of our acquired PPIs with the STRING database. Among the 3,366 PPIs, 1,185 were found in the STRING database and matched to the corresponding reliability score. As shown in Fig. 3B, 62% (729 PPIs) of our identified protein interactions were in the highest score range of 0.9-1.0, while more than 72% of all the human protein interaction data collected in the STRING database were in the relatively low score range of 0.1-0.3. Two reasons may primarily contribute to the high credibility of the protein interactions achieved by our method. First, more accurate interactions might be acquired from living cell experimental conditions. Second, the cross-linked interface site information could provide direct information on the interactions of the protein complexes. Taking interprotein cross-links of BIP-PDIA6 and BIP-HYOU1 as an example, BIP plays a role in facilitating the assembly of multimeric protein complexes inside the endoplasmic reticulum that are involved in the correct folding of proteins and degradation of misfolded proteins, and its interacting proteins PDIA6 and HYOU1 both function as chaperones that participate in protein folding and in inhibiting the aggregation of misfolded proteins. We identified two cross-links between PDIA6 and BIP and another two cross-links between HYOU1 and BIP, providing structural binding relevance that was complementary to the current evidence suggesting functional links. Taken together, these results demonstrated the great capability of our method in capturing protein interactions with different functions in various cellular localizations within living cells, further promoting our understanding of the underlying molecular mechanisms of cellular processes.

**Fig. 3.**
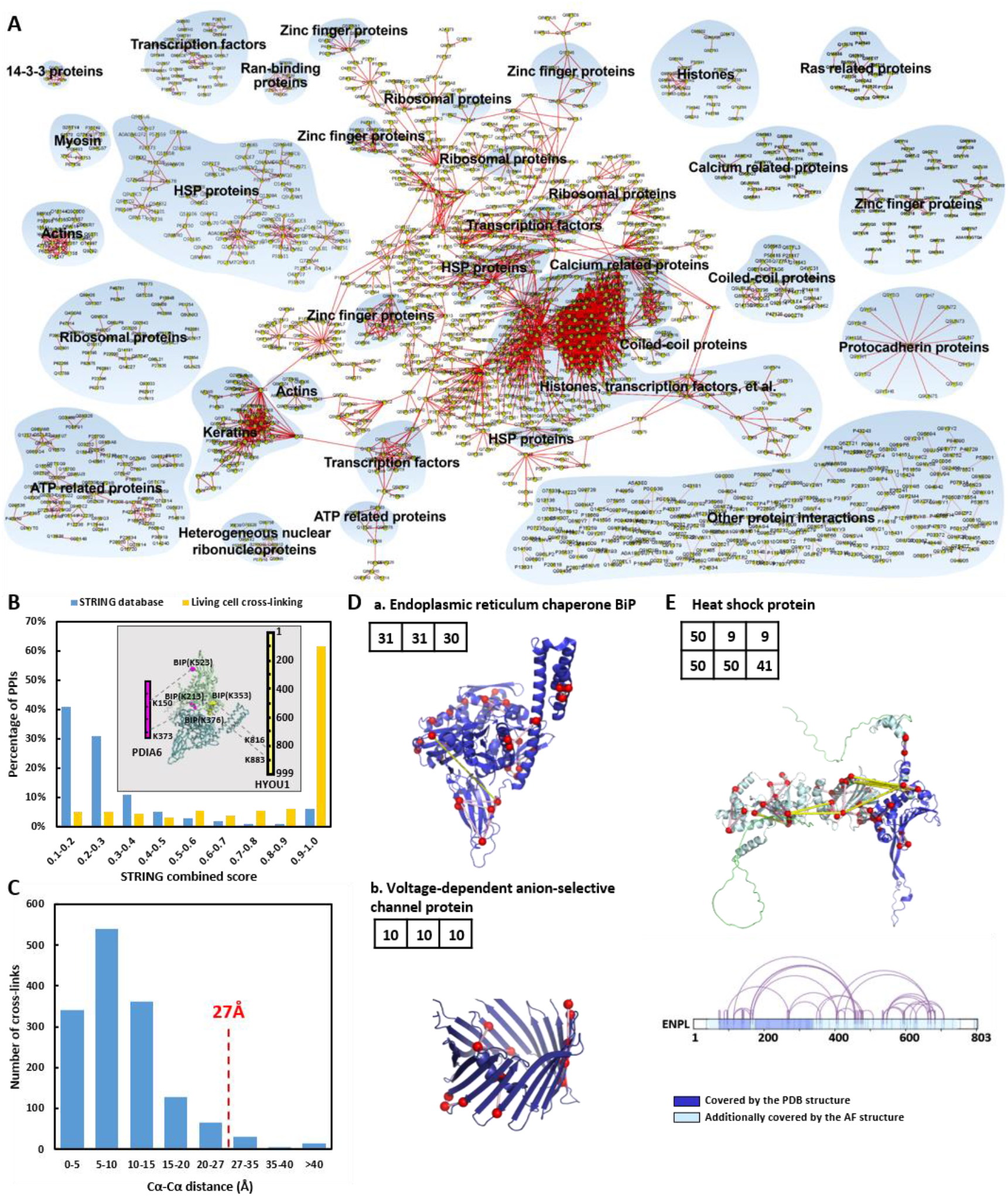
Validation of the interprotein PPIs and the intraprotein cross-links mapping to structural architecture. (*A*) The entire PPI network. This map consists of 1,595 nodes and 3,366 edges with simple biological process benchmarking. (*B*) Comparison of the STRING score distribution between our identified PPIs and all the human PPIs in the STRING database. The cross-linked interface of BIP-PDIA6 and BIP-HYOU1 is shown as an illustration. STRING score distributions for human PPIs in STRING (blue) and our PPIs (yellow). (*C*) Distance distribution of all the intraprotein cross-links mapped onto the available high-resolution structures in PDB. We used Cα–Cα Euclidean distances (ED) measured using the straight-line distance of the atomic coordinate in a PDB file. (*D*) a. Cross-links mapped on the structure of the endoplasmic reticulum chaperone BiP (PDB: 6ASY); b. Cross-links mapped on the structure of the voltage-dependent anion channel protein (PDB: 6TIR). Cross-links shown as pink lines could be mapped on the high-resolution structure within the distance constraint of 27 Å, while yellow lines exceeded 27 Å. The cross-linked Lys residues are shown as red spheres. The numbers on the upper left corner of the structure represent in turn the number of identified cross-links of the protein complex, the number of cross-links that could be mapped on the crystal structure, and the number of cross-links within the cross-linking constraint. (*E*) Cross-links mapped on the structure AF-P14625 of HSP90 from *AlphaFold2*. All the cross-links identified within the protein are shown using the linear connectivity map. The structural sequence covered by the experimentally determined structure is colored blue. Additional structural sequences covered by the *AlphaFold2* structure are colored pale cyan. The description of the top row data was the same as that in Fig. 3D. For the row below, the numbers represent in turn, the number of identified cross-links of the protein complex, the number of cross-links that could be mapped on the AF structure and the number of cross-links within the cross-linking constraint.

In addition, we identified 13,857 intraprotein cross-linked sites corresponding to 2,128 proteins. To evaluate cross-linking validity, we mapped all the cross-linked sites onto the available high-resolution structures of protein complexes in the Protein Data Bank (PDB). In total, 1,484 cross-linked sites could be mapped onto their respective structures among 543 proteins, with 97% satisfaction within the maximum Cα–Cα distance restraint of 27 Å (Fig. 3C), confirming the validity of our identified cross-links and suggesting that our cross-linking conditions did not significantly disturb protein structural conformations. Then, to test the universality of our data, some of the cross-links were mapped on representative protein structures with different cellular locations (Fig. 3D, *SI* Fig. S25, Dataset S4). Nearly all the cross-links could be mapped on the crystal structure and had high structural compatibility (from 84% to 100%). In addition, not all the identified cross-links could be mapped to the available crystal structure. For the heat shock protein HSP90, only the CATH domain (73-337) was resolved with X-ray. We identified 50 intraprotein cross-links of HSP90, 9 of which were within the CATH domain and could be mapped to the crystal structure with Cα-Cα distances completely satisfying the distance restraint. Then, we expanded the structural coverage by applying the recently reported highly accurate protein structure prediction tool *AlphaFold2* (AF) (32). The strong correlation of our identified cross-links and the AF predicted structure with 82% structural compatibility indicates that our method is a promising experimental tool for protein structure analysis (Fig. 3E, Dataset S4).

### Spatially resolved profiling of protein complex by biocompatible chemical cross-linking in living cells

Eukaryotic cells are highly compartmentalized with a unique microenvironment (33). Currently, approximately half of all human proteins are known to localize in multiple compartments to dictate distinct functions. Most proteins undergo conformational changes and interaction dynamics in different subcellular localizations during translocation. Inspired by the ability of our living cell cross-linking method to capture the assembly of protein complexes with high biocompatibility and deep coverage, it was further extended to profile spatially specific protein conformations and interactions (Fig. 1).

The tumor suppressor phosphatase and tensin homolog PTEN is a negative regulator of the PI3K/AKT pathway. Early studies (34) found that endogenous wild-type PTEN was expressed mainly in the cytoplasm, and a relatively small amount was exclusively localized in the nucleus. In addition, nuclear PTEN was dramatically increased in the presence of the proteasome inhibitor MG132 (35), which was also verified in our experiment (Fig. 4A, *SI* Note S12). In this work, we performed SPACX to explore the spatial heterogeneity of conformation and interaction for PTEN, as well as the PTEN nuclear translocation process (Fig. 4B). Primarily, high purity separation is of great significance. We performed living cell cross-linking coupled with efficient separation of the cytoplasm and nucleus (Fig. 4C). Briefly, 18 and 45 cross-linked sites within PTEN were identified in the cytoplasm and nucleus (Dataset S5), respectively. Mapping the cross-links to the reported PTEN crystal structure (PDB: 5BUG) revealed 4 and 6 cross-links between the phosphatase tensin-type domain and the C2 tensin-type domain in the cytoplasm and nucleus, respectively, exceeding the maximum distance constraint of BSPNO cross-linking. By ensemble structure refinement with cross-linking restraints (36) (*SI* Note S13), subcellular-specific conformations were presented for PTEN. The nucleus displayed a broader range of dynamic conformational changes in the dual specificity domains than the cytoplasm (Fig. 4D).

**Fig. 4.**
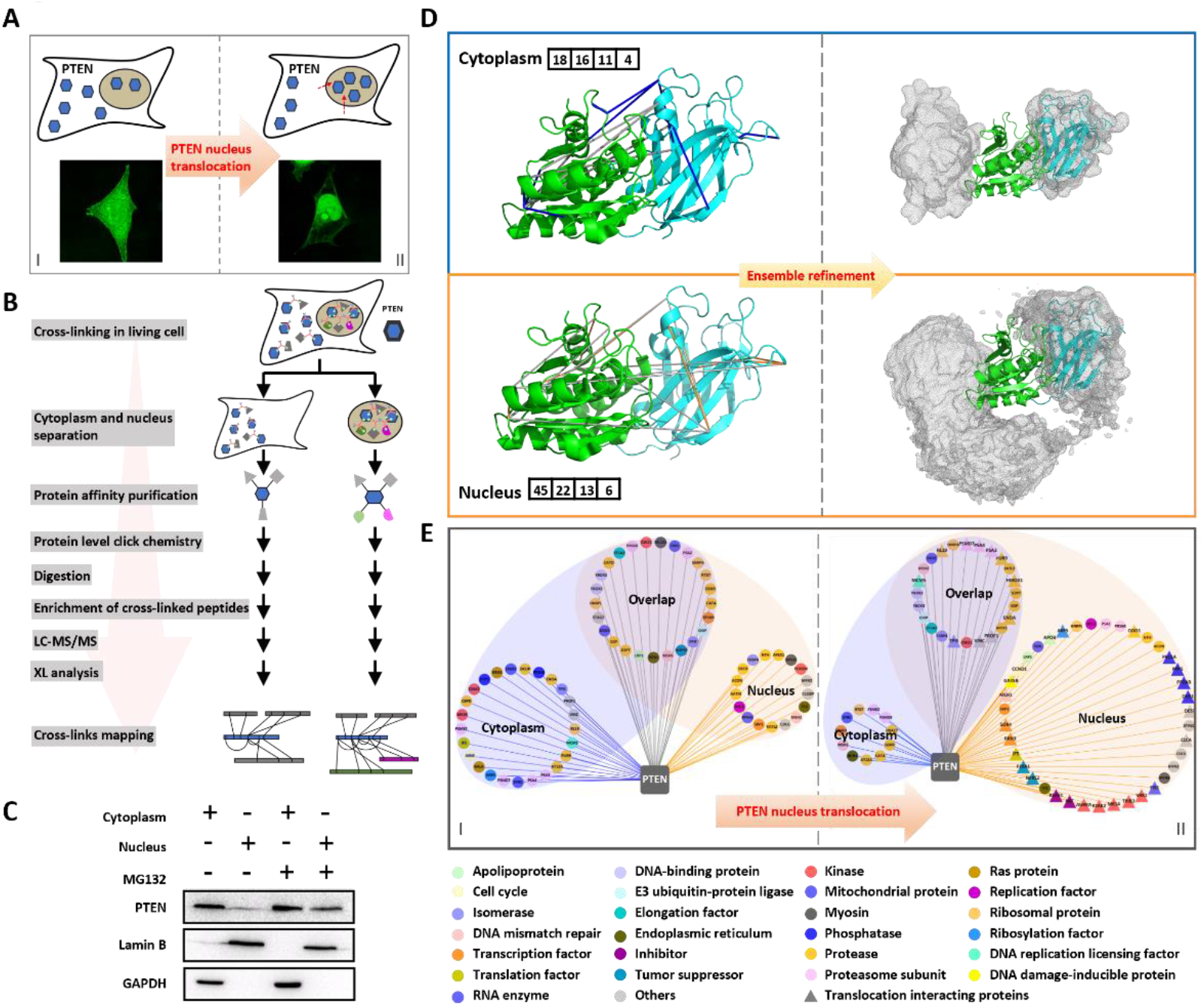
SPACX-enabled subcellular specific PTEN conformation and interactome analysis. (*A*) Diagram and fluorescence imaging analysis of PTEN localization and nuclear translocation. (*B*) Schematic description of SPACX for PTEN analysis. (*C*) Western blot analysis of the purity of the separation of the cytoplasm and nucleus and the diversity of PTEN expression levels. GAPDH and Lamin B served as cytoplasmic and nuclear markers, respectively. (*D*) Cross-links mapped on the PTEN structure PDB 5BUG and ensemble refinement for PTEN conformational changes against the cross-linking restraints. Upper, cytoplasm; lower, nucleus. Cross-links shown as blue/orange lines were mapped on the high-resolution structure within the distance constraint of 27 Å, while gray lines exceeded 27 Å. The green-colored structure was a phosphatase tensin-type domain, and the cyan-colored structure was a C2 tensin-type domain. The numbers on the side of the structure represent in turn the total number of cross-links; the number of cross-links mapped on the structure PDB 5BUG; the number of cross-links within the cross-linking constraint; and the number of cross-links used for ensemble refinement that extended between the two domains and exceeded the cross-linking constraint. (*E*) PTEN interaction networks. The proteins found to have direct interactions with PTEN are illustrated as individual nodes. The interacting proteins are color coded based on functional annotation. The interacting proteins shown as triangles were generated from PTEN nuclear translocation.

With such dynamic conformational differences, different interacting assemblies would form to exert diverse biological functions. Altogether, we identified 50 cytoplasmic and 42 nuclear PTEN-interacting proteins, including 25 overlapping interacting proteins (Fig. 4E, Dataset S6). In addition, the fluorescence colocalization and Western blotting results further certified the cross-linking results from cytoplasm- and nucleus-specific localization and the corresponding interactions (*SI* Fig. S26 and Note S14). In addition, for the exclusively identified interacting proteins in the cytoplasm, MK14 was enriched in the BAG2 signaling pathway; ENOA and TPIS in the glycolysis I pathway; PSA3, PSA4, PSME2 and PSMD7 in the protein ubiquitination pathway; ARF5, RALA, RRAS, VINC and PROF1 in the integrin signaling pathway; and E2AK2 in the PDGF signaling pathway, which were all closely related to the PI3K-AKT signaling pathway (*SI* Fig. S27). Among the nucleus-exclusively interacting proteins identified, MSH2, APEX1, NTH and RFC2 were mainly enriched in the DNA damage and repair pathways, which were distinct from those in the cytoplasm (*SI* Fig. S28). Therefore, all these results demonstrated the ability of our method to profile subcellular-specific protein complex assemblies, which exemplified the importance of SPACX for exploring protein structural and functional heterogeneity in diverse intracellular locations.

Finally, we extended SPACX to monitor the dynamic interaction changes involved in PTEN translocation from the cytoplasm to the nucleus upon ubiquitin-proteasome system (UPS) stress (35). By comparison, 36 PTEN-interacting proteins were newly produced within the nucleus due to PTEN nuclear translocation. All these proteins were identified with higher intensity in the nucleus after stimulation (Dataset S6). Among them, 6 proteins were completely moved from the cytoplasm to the nucleus after stimulation; 10 proteins were found to interact with PTEN partially in the nucleus after stimulation; and 20 proteins were newly found to interact with PTEN after stimulation (Fig. 4E). Five of the newly generated PTEN-interacting proteins were verified by Western blot and fluorescence colocalization analysis. PPP5, MK14 and PP2AB were found to interact with PTEN in the cytoplasm before stimulation and in the nucleus after stimulation; PROF1 was cross-linked with PTEN in the cytoplasm before stimulation and in both the cytoplasm and nucleus after stimulation; and RRN3 was found to interact with PTEN only after stimulation. The Western blot and fluorescence colocalization results corroborated the cross-linking results (Fig. 5).

**Fig. 5.**
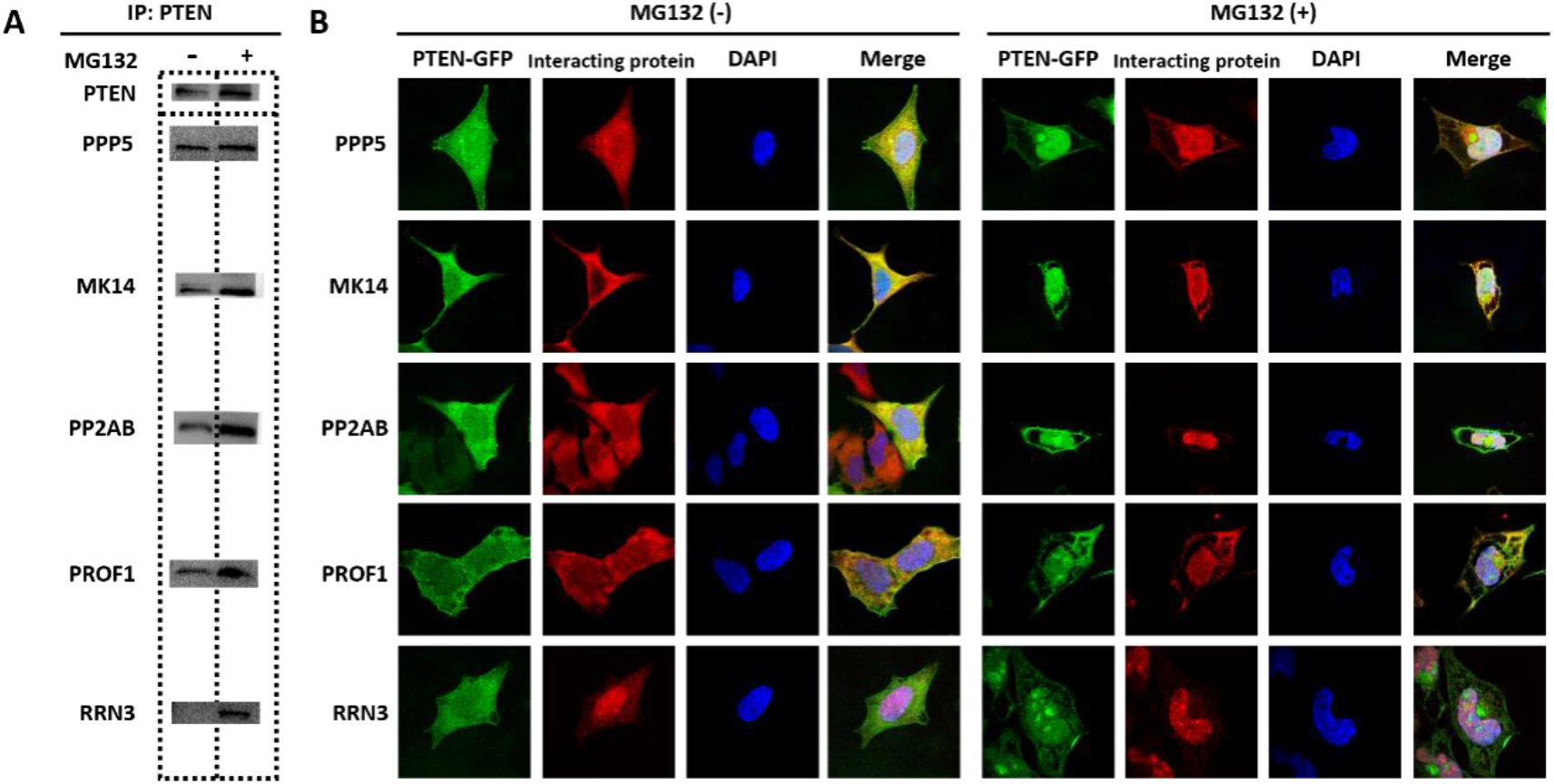
Validation of PTEN nuclear translocation interacting proteins. (*A*) Western blot analysis of the PTEN-interacting proteins PPP5, MK14, PP2AB, PROF1 and RRN3 by pulling down PTEN. (*B*) Immunofluorescence imaging analysis of the colocalization of PTEN (green, GFP) and the PTEN nuclear translocation interacting proteins PPP5, MK14, PP2AB, PROF1 and RRN3 (red, Alexa Fluor 647-conjugated antibody). Merge includes demarcated nucleus (blue, DAPI).

Then, Ingenuity pathway analysis (IPA) was used to enrich the functional pathways involved in the newly generated nucleus PTEN-interacting proteins. CCND1, E2AK2, TRIB3, PR15A and RL19 were enriched in the EIF2 signaling pathway; MK14, PP2AB and GA45B in the ATM signaling pathway; CCND1, MK14 and GA45B in the p53 signaling pathway; and all were directly related to cell apoptosis (*SI* Fig. S29), indicating that PTEN nuclear translocation might induce cell apoptosis upon UPS stress treatment.

In addition, PTEN exists in three isoforms with distinct functions. Identifying individual protein isoforms and interacting proteins is challenging but attractive for understanding their functional heterogeneity. Herein, we investigated the interactions between PTEN isoforms and their interacting proteins within the nucleus after stimulation. Taking isoform1 as the canonical sequence, isoform2 was produced by alternative initiation at a CTG start codon of isoform1, and isoform3 was produced by alternative splicing of isoform1 (*SI* Fig. S30). In total, we identified 7 cross-links between the unique sequence region (USR) of isoform2 and isoform1 and 20 cross-links between the USR of isoform3 and isoform1 (Fig. 6A, Dataset S7). The cross-links between PTEN isoforms could provide reliable interaction interfaces for modulating the direct interactions.

**Fig. 6.**
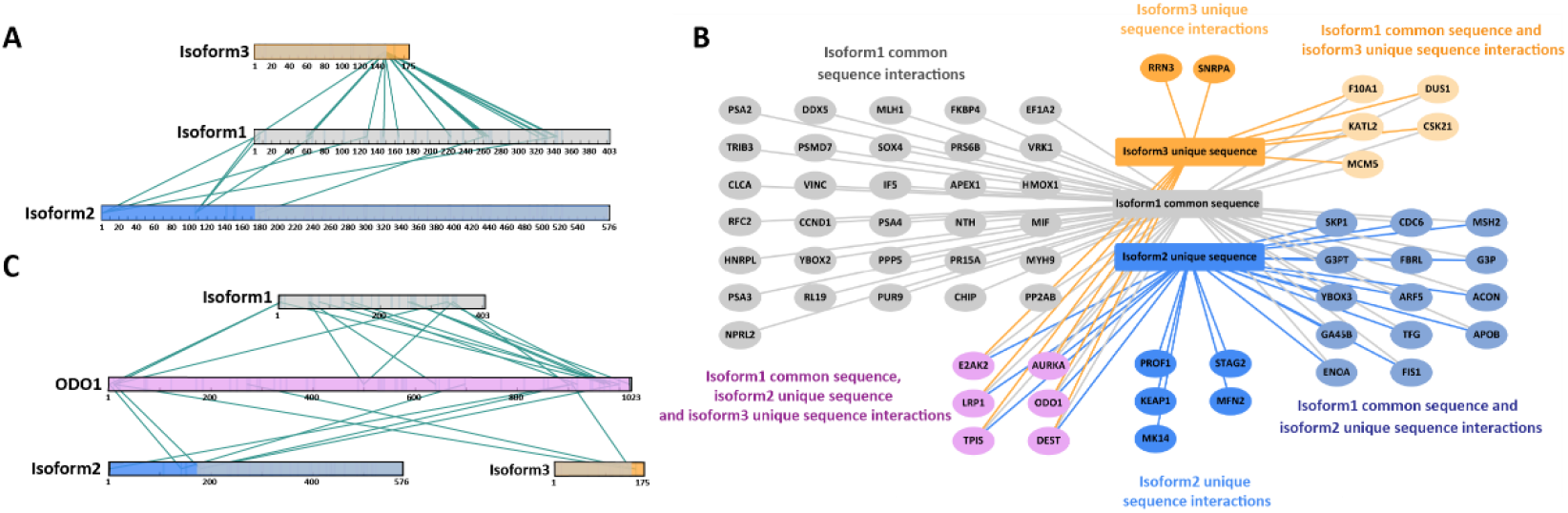
Analysis of PTEN isoform-containing assemblies. (*A*) Cross-links within homologous oligomers formed between PTEN isoforms are shown using linear connectivity maps. (*B*) PTEN isoform-containing interactions in the nucleus after UPS stimulation. The interacting proteins were color coded based on the interaction with the respective PTEN isoforms sequences. The color of the edges indicates which sequence region was cross-linked to the protein. (*C*) The cross-links between ODO1 and PTEN isoforms are shown using linear connectivity maps.

In addition to the acquisition of PTEN homodimers, we found all three PTEN isoforms interacting with 63 proteins. Among them, 5 proteins exclusively interacted with isoform2 USR, and 2 proteins interacted only with isoform3 USR. All the other proteins interacted with multiple PTEN isoforms (Fig. 6B). The protein 2-oxoglutarate dehydrogenase ODO1, which was exclusively identified in the nucleus after stimulation, was found to interact with all three PTEN isoforms (Fig. 6C, Dataset S7). Multiple cross-links were obtained to provide reliable interaction interfaces for direct interaction determination. ODO1 is mainly active in the mitochondrion and is required to localize in the nucleus as well for the lysine succinylation of histones (37, 38). Our findings might allow further understanding of PTEN isoform interactions in the cell cycle and transcription regulation. Taken together, these results demonstrated the capability of our SPACX strategy in characterizing the dynamic changes of protein conformations and interactome during subcellular translocation in living cells for biological function exploration, especially in the functional specificity of protein isoforms.

## Discussion

Understanding the functional mechanisms of proteins during cellular biological processes requires comprehensive analysis of subcellular specific protein conformation changes and interaction dynamics. In this work, we developed the SPACX strategy by chemical cross-linking in living cells followed by subcellular separation to characterize protein conformational and interactional dynamics. First, based on the fast membrane permeability, high reactivity and good water solubility of the cross-linker BSPNO, as well as the constructed biocompatible cross-linking conditions, cross-linking was possible with minimal intrusion on cell viability and protein stability in living cells. In addition, bioorthogonal enrichment of cross-linked peptides by introducing a biotin tag via alkyne-azido click chemistry allowed high identification sensitivity and deep coverage. The advancement of in vivo capture of protein complexes in physiological environments within 5 min, together with the highly efficient enrichment and identification of cross-links, enhanced our detection of the dynamic assembly of protein complexes in living cells on the global scale. This was a prerequisite to decipher the detailed dynamic change of spatial protein conformations and interactome.

Briefly, 18,886 interprotein cross-linked sites were confidently identified in human BEL7402 cells and mapped to 3,366 PPIs involving 1,595 proteins and 13,857 intraprotein cross-linked sites among 2,128 protein structural constraints, associated with diverse functions in almost all cellular compartments. Notably, our dataset comprehensively included some highly dynamic and weak binding interactions, such as histones, HSPs and TFs, that were hardly detectable by in vitro methods. This superiority was mainly contributed by the fast capture of cross-linking and the high protein concentration arising from the crowding effect in living cells, which also permitted the capability to map flexible structural regions that are difficult to capture in crystal structures but are important in protein recruitment and interaction formation.

Subsequently, by integrating with highly purified subcellular separation, spatially resolved characterization of the conformations and interactions of protein complexes within subcellular specific locations was available. By structural refinement against the cross-linking restraints, PTEN conformation revealed a broader range of dynamic changes in the dual specificity domains in the nucleus than in the cytoplasm. In addition, diverse interacting networks were found based on the differential conformations with our SPACX method. Through biological function analysis, separate functions of PTEN-interacting proteins were acquired in the cytoplasm and nucleus, enabling the exploration of the spatially specific structure, interaction and function of protein assemblies.

Finally, we demonstrated the feasibility of SPACX in studying the dynamic changes of interaction involved in protein translocation between subcellular spaces. The validity of interaction and colocalization for the newly produced PTEN-interacting proteins in nuclear import was certified by Western blot and fluorescence colocalization analysis. In addition, the homologous oligomers and individual interactions of PTEN isoforms were accessible. Therefore, our SPACX method could provide a general toolkit for the spatially resolved profiling of dynamic protein conformation and interactome in living cells.

