## supporting information for "Spatially resolved profiling of protein conformation and interactions by biocompatible chemical cross-linking in living cells"

[a]CAS Key Laboratory of Separation Science for Analytical Chemistry, National Chromatographic R. & A. Center, Dalian Institute of Chemical Physics, Chinese Academy of Sciences, Dalian, Liaoning 116023, China; [b]University of Chinese Academy of Sciences, Beijing 100039, China; [c]Innovation Academy for Precision Measurement Science and Technology, Chinese Academy of Sciences, Wuhan 430071, China; [d]Beijing National Laboratory for Molecular Sciences, College of Chemistry and Molecular Engineering, Beijing 100871, China; [e]Peking-Tsinghua Center for Life Sciences, Peking University, Beijing 100871, China

National Chromatography R. & A. Center
Dalian Institute of Chemical Physics
Chinese Academy of Sciences
Dalian 116023, China

### SUPPLEMENTARY METHODS

**Note S1 - Synthesis of BSPNO.**

The cross-linker BSPNO was synthesized according to a previous report (1) with minor modifications by three reaction routes. Briefly, first compound 1 was obtained by the acidification of tris (2-cyanoethyl) nitromethane in hydrochloric acid. Then, the three carboxyl groups were activated by forming NHS esters to produce compound 2. Next, BSPNO was prepared by the selective amidation of compound 2 with propargylamine. Finally, the target product BSPNO was purified by semipreparative HPLC. Detailed synthetic procedures and compound characterization are given in the supplementary data below.

**Note S2 - Calculation of the maximum Cα–Cα distance restraint of BSPNO.**

For the BSPNO arm length calculation, we used the molecular dynamics option of Chem3D 19.0 with standard settings applied (Step interval = 2.0 fs, Frame interval = 10 fs, Terminate After = 10000 steps, Heating/Cooling Rate = 1.000 Kcal/atom/ps, and Target Temperature = 300 K). The maximum Cα-Cα distance of BSPNO is an arm length of 8.9 Å, adding two lysine side chain lengths of 12 Å and a typical tolerance of 6 Å, which is close to 27 Å.

**Note S3 - Calculation methods of the surface area and cLogP of the cross-linkers.**

We calculated the surface area in PyMOL (function 'get_area'). We estimated cLogP by calculation with ChemDraw Professional 19.0.

**Note S4 - Experimental procedure of CCK-8 measurement.**

HeLa cells were seeded at 5000 cells per well in 96-well plates. The cells were washed three times with 1× PBS and cross-linked with 2 mM BSPNO (1× PBS, 1% DMSO (v/v)) for different time and then washed three times with 1× PBS again. Subsequently, 100 μL of MEM and 10 μL of CCK-8 reagent were added to the cells. The cells were incubated at 37 °C for 2 h. Optical density was measured at 450 nm using a MULTISKAN GO (Thermo Scientific). Six replicates were prepared for each cross-linking time.

**Note S5 - Experimental procedure of cell morphology observation.**

The cells were cultured in the dish, washed three times with 1× PBS and then cross-linked with 2 mM BSPNO (1× PBS, 1% DMSO (v/v)). Images of the same view were taken at different cross-linking time with a microscope (Nikon ECLIPSE Ti-S) and analyzed using Nikon NIS-Elements D.

**Note S6 - Data processing for label-free quantitation.**

The label-free quantitation raw files were processed with MaxQuant (v1.5.1.0), supported by the Andromeda search engine. The raw data were searched against the *Homo sapiens* fasta. Carbamidomethyl[C] was set as fixed modification. Acetyl[Protein N-term], Oxidation[M], BSPNO-COOH (+296.1008), BSPNO-NHS (+393.1172) and BSPNO-NH2 (+295.1168) were set as variable modifications. BSPNO was assumed to react with lysine and the protein N-terminus. The other parameters were set up as follows: full tryptic specificity, up to three missed cleavages; label-free quantification, LFQ; match between runs, match time window set as 0.7 min, alignment time window set as 90 min; use both unmodified peptides and the peptides with the above variable modifications for quantitation; the FDR was set to 0.01 both at the protein and the peptide spectra match (PSM) level. The database on *Homo sapiens* (42,432 reviewed proteins, including canonical and isoform) was downloaded from UniProt on 2019-04-04.

**Note S7 - Cross-linking of BSA with BSPNO for different time.**

BSA was dissolved in the cross-linking buffer (20 mM HEPES, 150 mM NaCl, pH 7.8) and cross-linked with 2 mM BSPNO at room temperature for 3 min, 5 min, 10 min, 15 min, 30 min and 60 min. Next, the cross-linking reaction was quenched with 40 mM ABC. Then, the proteins were deposited by acetone precipitation, and the precipitated protein pellets were air dried and resuspended in 8 M urea (50 mM ABC). Following by reduction (10 mM TCEP, 25 °C, 1 h) and alkylation (20 mM IAA, 25 °C, 20 min, dark), the samples were diluted to 1 M urea with 50 mM ABC and digested with trypsin at 37 °C overnight.

**Note S8 - Fluorescence imaging characterization of BSPNO cell membrane permeability.**

HeLa cells were cultured in MEM in 35 mm Petri dishes with a number 1.5 cover glass bottom until they reached 70%-80% confluence. The cells were washed five times with 1× PBS and then treated with 2 mM BSPNO (in 1× PBS containing 1% DMSO) for 5 min at room temperature. Following the cross-linking reaction, the cells were washed 5 times with 1× PBS and fixed with 10% HCHO, 1× PBS for 15 min at room temperature with constant shaking. After fixation, the cells were incubated with 0.1% Triton X-100 in 1× PBS for 30 min for membrane perforation. The click-chemistry reaction was performed with 10 mM biotin-PEG3-$N_3$ in a mixture of $CuSO_4$, sodium ascorbate and THPTA ligand for 2 h at room temperature with constant shaking. After washing with 1× PBS, the cells were incubated with 1 μg/mL Streptavidin Fluor 488 (Thermo Fisher) for 1 h in

0.1% Triton X-100, 1× PBS buffer and 1 μg/mL DAPI in 1× PBS for 15 min in the dark. The cells were then washed with 1× PBS for 5 min in the dark. The control group cells were treated with 1% DMSO in 1× PBS for 5 min in the cross-linking step, which was the only difference from the experimental group. Confocal fluorescence imaging was performed separately at the 365 nm and 488 nm channels using a 100 × objective.

**Note S9 - Experimental procedure of cross-linked peptides enrichment.**

The cross-linked proteins were dissolved in 0.2% SDS (1x PBS). The same volumes of 160 mM THPTA, 20 mM $CuSO_4$ and 500 mM sodium ascorbate were added to the protein solution. ADB enrichment reagent (DMSO, 20 mM) was added at a 2:1 volume ratio of 160 mM THPTA and allowed to react at 60 °C for 2 h. After the click-chemistry reaction, the proteins were deposited by acetone precipitation and the precipitated protein pellets were air dried and resuspended in 8 M urea (50 mM ABC). Following by reduction (5 mM TCEP, 25 °C, 1 h) and alkylation (10 mM IAA, 25 °C, 20 min), the samples were diluted to 1 M urea with 50 mM ABC and digested with trypsin at 37 °C overnight. Subsequently, the peptides were incubated with streptavidin beads at room temperature for 1 h, and then the beads were washed with 1 M KCl (1x PBS), HPLC $H_2O$ and 10% ACN (90% $H_2O$) to remove nonspecific absorption. Finally, the cross-linked peptides were released from the beads with $Na_2S_2O_4$ in 6 M urea and 2 M thiourea buffer.

**Note S10 - Experimental procedure of *E. coli* lysate proteins cross-linking.**

The *E. coli* cell suspensions were harvested at OD 0.6-0.8. The cells were pelleted and washed 3 times with 1× PBS, and then twice with 20 mM HEPES containing 50 mM NaCl and 1.5 mM $MgCl_2$, pH 7.4. Then, the sample was added to lysis buffer (20 mM HEPES, 50 mM NaCl, 1.5 mM $MgCl_2$, 1% cocktail, pH 7.4), ultrasonicated on ice, and centrifuged at 16,000 ×g for 30 min to remove the deposit. The protein concentration was determined with the BCA assay. The cross-linking reaction was performed with a weight ratio of protein to cross-linker of 2:1 (BSPNO) and 4:1 (DSS) in lysis buffer at room temperature for 1 h. After that, the cross-linking reaction was quenched with 50 mM ABC.

**Note S11. Cross-linking of BEL7402 cell lysate proteins with BSPNO.**

Cells were harvested by trypsinization and washed three times with 1× PBS. The cell pellet was first lysed in cellular membrane lysis buffer (10 mM HEPES, 10 mM KCl, 1.5 mM $MgCl_2$, 0.5 mM DTT, 0.4% NP-40, pH 7.8, 1% cocktail) for 10 min on ice. Then, the nuclei were pelleted by centrifugation for 10 min at 3,200 ×g. The nuclear pellet was resuspended in cross-linking buffer (20 mM HEPES, 150 mM NaCl, 1.5 mM $MgCl_2$, 0.5 mM DTT, pH 7.8, 1% cocktail) and lysed by sonication on ice for three cycles (30 s in each cycle with 5 s on, 20 s off) with 50% amplitude power. The supernatant fraction was cytoplasmic proteins. The protein concentration was determined with the BCA assay. The cross-linking reaction was performed with a weight ratio of protein (2 mg/mL) to cross-linker (BSPNO) of 4:1 at room temperature for 1 h and then quenched with 50 mM ABC.

**Note S12 – UPS stress stimulus condition.**

293A PTEN-GFP-biotin cells were treated with 10 μM MG132 for 24 h.

**Note S13 - Ensemble refinement of PTEN against the cross-linking restraints.**

We first analyzed the distance of the cross-links on the crystal structure of PTEN and found that many cross-links exceeded the maximum distance constraint of the cross-linker. Thus, ensemble refinement against the cross-linking restraints was performed using Xplor-NIH. The ensemble comprised two or more conformations that collectively accounted for the cross-linking results. Ambiguous distance restraints were used for the cross-links in the nucleus and cytoplasm. The distance between Cβ atoms was used as the distance restraint. The maximum distance was set to 20 Å according to the arm length of the cross-linker. The lowest limit of 4 Å was set according to the van der Waals radius of the atoms. The square-well energy function was used, and the balance position was set to 10 Å from the maximum probability of distance distribution in molecular dynamics simulations.

The crystal structure of PTEN (PDB: 5BUG) was used for structure refinement. In addition to the distance restraints, other knowledge-based potentials, including covalent energy (bond, angle) and van der Waals, were applied to the entire protein. During ensemble refinement, each cross-linking restraint was accounted for with either the ground state (crystal structure) or an excited state with an alternative domain arrangement. In each conformer, the N-terminal domain (residues 14-184) was fixed, and the C-terminal domain (residues 193-327) was grouped and moved together as a whole. The linker residues between the N-terminal domain and C-terminal domain (residues 185-192) had full torsion angle freedom. The conformers in the ensemble shared all the cross-linking restraints and were allowed to overlap. If the two-conformer ensemble could not satisfy all the cross-linking restraints, three or more conformers were added to the ensemble until all the restraints are satisfied.

We performed the calculation 960 times for PTEN with different random seeds. Structures with no violation of cross-linking restraints and no clashed residues were used for further structural analysis. The atomic probability maps depicting the distribution of the C-terminal domain relative to the N-terminal domain were calculated in Xplor-NIH and plotted at the corresponding thresholds. The structure figures were rendered using PyMOL (version 2.2, Schrödinger).

**Note S14 - Western blot and immunofluorescence analysis.**

Antibodies used for WB analysis, anti-PTEN (1:1000), anti-GAPDH (1:10000), anti-Lamin B (1:10000), anti-NTH (1:1000), anti-RRN3 (1:1000), anti-PROF1 (1:10000), anti-CCND1 (1:1000), anti-IF5 (1:1000), anti-PPP5 (1:1000), anti-MK14 (1:1000), anti-PP2AB (1:1000), HRP-conjugated anti-rabbit IgG (H+L) Secondary Antibody (1:20000).

Antibodies used for immunofluorescence analysis include anti-NTH (1:150), anti-RRN3 (1:100), anti-PSME2 (1:500), anti-PROF1 (1:100), anti-CCND1 (1:100), anti-IF5 (1:100), anti-PPP5 (1:100), anti-MK14 (1:100), anti-PP2AB (1:100) and anti-E2AK2 (1:100).

**Note S15 - Peptide sample fractionation before MS analysis.**

The living cell cross-linked BEL7402 cells samples and PTEN nucleus fraction samples were respectively desalted and fractionated with home-made Durashell C18 Tips. Mobile phase A was $H_2O$ adjusted to pH 10 using $NH_3 \cdot H_2O$ and mobile phase B was 80% ACN and 20% $H_2O$ adjusted to pH 10 using $NH_3 \cdot H_2O$. Samples were in order eluted with 6% ACN, 9% ACN, 12% ACN, 15% ACN, 18% ACN, 21% ACN, 25% ACN, 30% ACN and 80% ACN. The flow through of 6% and 25% ACN were made fraction 1, 9% and 30% ACN were made fraction 2, and 12% and 80% ACN were merged into fraction 3. The eluent from 15%, 18% and 21% ACN were used as fraction 4, 5 and 6, respectively.

PTEN cytoplasm fraction samples were also desalted and fractionated with home-made Durashell C18 Tips. Mobile phase composition and elution order were the same with nucleus fraction. While, the flow through of 6%, 15% and 25% ACN were made fraction 1, 9%, 18% and 30% ACN were made fraction 2, and 12%, 21% and 80% ACN were merged into fraction 3.

BSPNO cross-linked BEL7402 cell lysate samples and BSPNO cross-linked BSA samples were desalted with home-made Venusil XBP C18 Tips. Mobile phase A was $H_2O$ with 0.1% TFA and mobile phase B was 80% ACN, 20% $H_2O$ with 0.1% TFA. Samples were desalted with mobile phase A and eluted with mobile phase B.

Fractionation of BSPNO cross-linked BEL7402 cell lysate peptides before enrichment. The peptides were fractionated by high pH RPLC with a C18 column (250 mm×4.6 mm, Durashell). Mobile phase A (98% $H_2O$, 2% ACN, adjusted pH to 10 using $NH_3 \cdot H_2O$) and B (98% ACN, 2% $H_2O$, adjusted pH to 10 using $NH_3 \cdot H_2O$) were used to set the gradient as follows: 4%-45% B, 45 min; 45%-100% B, 5 min; 100% B, 50 min. The flow rate was 1 mL/min. The eluent was collected every minute. A total of 50 fractions were collected, and then concatenated to 10 fractions with equal time interval.

**Note S16 - LC-MS/MS analysis.**

The label-free quantification samples, BSPNO cross-linked BEL7402 cells samples, BSPNO cross-linked BEL7402 cell lysate samples and BSPNO cross-linked PTEN protein complex samples were all analyzed with an Easy-nLC 1200 system coupled to an Orbitrap Fusion Lumos mass spectrometer. Mobile phase A consisted of 0.1% FA in HPLC $H_2O$, and mobile phase B of 80% ACN, 0.1% FA and 20% HPLC $H_2O$.

The label-free quantification samples were analyzed using a 90 min gradient with the flow rate of 600 nL/min. The gradient was set as follows, 62 min from 5% to 35% B, 21 min from 35% to 50% B, 2 min from 50% to 95% B and 95% B maintained for 5 min. The mass spectrometry was operated in data-dependent mode with one full MS scan at R = 60,000 (m/z = 200), followed by MS/MS scans at R = 15,000 (m/z = 200), RF Lens (%) = 30, with an isolation width of 1.6 m/z. The AGC target for the MS1 and MS2 scan were 400,000 and 50,000 respectively, and the maximum injection time for MS1 and MS2 were 50 ms and 30 ms. The precursors with charge states 2 to 7 with an intensity higher than 20,000 were selected for HCD fragmentation, and the dynamic exclusion was set to 20 s. Each sample was analyzed by three parallel runs and the TIC intensity was controlled to ensure the approximately same loading amount of samples.

The samples from BSPNO cross-linked cells were analyzed using a 75 min gradient with the flow rate of 600 nL/min (56 min from 5% to 34% B, 15 min from 34% to 55% B, 1 min from 55% to 95% B and 3 min 95% B). The precursors with charge states from 3 to 7 were selected for MS2 fragmentation and the other parameters were the same as label-free samples.

The samples from BSPNO cross-linked cell lysate were analyzed using a 75 min gradient with the flow rate of 600 nL/min (51 min from 4% to 25% B, 20 min from 25% to 50% B, 1 min from 50% to 95% B and 3 min 95% B). The MS parameters were the same as living cell cross-linked samples.

The remaining supernatant after BSPNO cross-linked cell lysate enrichment were also identified with LC-MS/MS to create a target database for the cross-linked peptides. The samples were analyzed with an Easy-nLC 1200 system coupled to an Orbitrap Fusion Lumos mass spectrometer using the same gradient with the cross-linked peptides. The only difference of the mass spectrometry parameters with the label-free quantification samples was MS2 detector type, iontrap, scan rate rapid, maximum injection time 15 ms and AGC target 10,000.

Different fractions of BSPNO cross-linked PTEN protein complex samples were all analyzed using an 85 min gradient with the flow rate of 600 nL/min, while the detail gradients were different. Cytoplasm fraction 1: 28 min from 6% to 16% B, 29 min from 16% to 34% B, 17 min from 34% to 48% B, 1 min from 48% to 95% B and 10 min 95% B. Cytoplasm fraction 2: 26 min from 10% to 16% B, 30 min from 16% to 34% B, 18 min from 34% to 48% B, 1 min from 48% to 95% B and 10 min 95% B. Cytoplasm fraction 3: 24 min from 12% to 16% B, 31 min from 16% to 34% B, 19 min from 34% to 48% B, 1 min from 48% to 95% B and 10 min 95% B. All the fractions combined cytoplasm sample: the same as cytoplasm fraction 3. Nucleus fraction 1: 18 min from 6% to 15% B, 39 min from 15% to 30% B, 17 min from 30% to 40% B, 1 min from 40% to 95% B and 10 min 95% B. Nucleus fraction 2: 8 min from 7% to 10% B, 35 min from 10% to 19% B, 2 min from 19% to 20% B, 29 min from 20% to 40% B, 1 min from 40% to 95% B and 10 min 95% B. Nucleus fraction 3: 17 min from 10% to 16% B, 23 min from 16% to 20% B, 30 min from 20% to 27% B, 5 min from 27% to 44% B, 1 min from 44% to 95% B and 9 min 95% B. Nucleus fraction 4: 5 min from 10% to 13% B, 17 min from 13% to 18% B, 41 min from 18% to 26% B, 12 min from 26% to 44% B, 1 min from 44% to 95% B and 9 min 95% B. Nucleus fraction 5: 1 min from 6% to 15% B, 16 min from 15% to 22% B, 54 min from 22% to 31% B, 3 min from 31% to 44% B, 1 min from 44% to 95% B and 10 min 95% B. Nucleus fraction 6: 1 min from 6% to 19% B, 23 min from 19% to 25% B, 44 min from 25% to 35% B, 6 min from 35% to 44% B, 1 min from 44% to 95% B and 10 min 95% B. All the fractions combined nucleus sample: 28 min from 6% to 16% B, 28 min from 16% to 34% B, 16 min from 34% to 48% B, 1 min from 48% to 95% B and 12 min 95% B. The MS parameters were the same as living cell cross-linked samples.

The cross-linked *E. coli* lysate samples were analyzed with an Ultimate 3000 RPLC nano coupled to an Orbitrap Velos mass spectrometer. Mobile phase A consisted of 0.1% FA in 98% HPLC $H_2O$ and 2% ACN, and mobile phase B of 98% ACN, 2% HPLC $H_2O$ and 0.1% FA. The samples were analyzed using a 108 min gradient with the flow rate of 600 nL/min (84 min from 6% to 28% B, 16 min from 28% to 40% B, 4 min from 40% to 95% B and 4 min 95% B). The mass spectrometer instrument was operated in positive mode with a 1.8 kV applied spray voltage. The temperature of the ion transfer capillary was set at 320 °C. One microscan was set for each MS and MS/MS scan. All MS and MS/MS spectra were acquired in the data dependent mode. A full scan MS acquired from m/z 350 to 1850 with charge states of 3-5, followed by 10 data dependent MS/MS events. The dynamic exclusion function was set as follows: repeat count, 1; repeat duration, 10 s; exclusion duration, 22 s. The normalized collision energy for MS/MS scanning was 40%.

The BSPNO cross-linked BSA samples were analyzed with an Easy-nLC 1000 system coupled to a Q-Exactive mass spectrometer. Mobile phase A consisted of 0.1% FA in 98% HPLC $H_2O$ and 2% ACN, and mobile phase B of 98% ACN, 2% HPLC $H_2O$ and 0.1% FA. The separation gradient was achieved by applying 2−7% B for 10 s, 7−23% B for 50 min, 23−40% B for 20 min, 40−80% B for 2 min, and 80% B for 13 min. The mass spectrometry was operated in data-dependent mode. The full MS scans were performed by the Orbitrap at 70,000@200 resolving power within the scan range of 300−1800 m/z. The AGC target for the full scans was 3000,000, and the maximum injection time was 60 ms. The loop count was 20, with the isolation window of 1.6 m/z. MS/MS scans were detected at the resolution of 17,500@200 with the fixed first mass of 110 m/z. The AGC target for the MS2 was 50,000, and the maximum injection time was 60 ms. Precursors were fragmented by higher-energy collision dissociation (HCD) with the normalized collision energy of 35%. Only precursors with charge states 3−7 with an intensity higher than 1000 were selected for fragmentation, and the dynamic exclusion was set to 20 s.

#### SUPPLEMENTARY DATASETS

**Dataset S1A.** Intraprotein cross-linked sites identified by our in vivo cross-linking.

**Dataset S1B.** Interprotein cross-linked sites identified by our in vivo cross-linking.

**Dataset S2.** Our identified interprotein interactions and overlap with the integrated PPIs databases of STRING and BioGRID and the selected published human CXMS studies.

**Dataset S3A.** Our identified histone protein interactions from in vivo cross-linking.

**Dataset S3B.** Our identified HSPs interactions from in vivo cross-linking.

**Dataset S3C.** Our identified TFs interactions from in vivo cross-linking.

**Dataset S4.** Detailed summary information of the Cα-Cα distance of the cross-links mapped to the corresponding PDB structures on Fig. 3 and SI Fig. 25.

**Dataset S5.** The Cα-Cα distance of intraprotein cross-links of PTEN in the cytoplasm and nucleus mapped to the PDB structure.

**Dataset S6.** PTEN interacting proteins in the cytoplasm and nucleus, with or without MG132 stimulation. We annotated protein function, cellular location and AP-MS intensity.

**Dataset S7.** Detailed cross-links information between PTEN isoforms, and ODO1 and PTEN isoforms.

### SUPPLEMENTARY FIGURES

**Supplementary Fig. S1 - Structural formula of BSPNO.**

**Supplementary Fig. S2 - Synthetic process of BSPNO.**

**Supplementary Fig. S3 - $^{1}$H NMR spectra of compound 1.**

**Supplementary Fig. S4 - MS spectra of compound 1.**

**Supplementary Fig. S5 - $^{1}$H NMR spectra of compound 2.**

**Supplementary Fig. S6 - MS spectra of compound 2.**

**Supplementary Fig. S7 - $^{1}$H NMR spectra of BSPNO.**

**Supplementary Fig. S8 - MS spectra of BSPNO.**

**Supplementary Fig. S9 - Chromatogram and separation condition of BSPNO.**

Purity was 99%, and retention time was 26 min, as detected by HPLC (A: $H_2O$, B: $CH_3CN$, gradient method as follows: from 5% B to 50% B over 40 min, 80% B over 10 min, 5% B over 10 min at a flow rate of 1 mL/min, monitored by UV wavelength at 200 nm).

**Supplementary Fig. S10 - Schematic diagram to assess the influence of cross-linking time on cell morphology, cell viability, protein structural stability and proteomic quantitative differences.**

**Supplementary Fig. S11 - Cell viability for different BSPNO cross-linking time.**

Cell viability was assessed for different BSPNO cross-linking time (0 min, 3 min, 5 min, 10 min, 15 min, 30 min, 60 min) measured by a CCK-8 kit. Values are represented as the mean ± s.e.m. All p values were > 0.05 from 3 min to 60 min vs. 0 min cross-linking (two-sided T-test), which was considered not statistically significant for all the tests.

**Supplementary Fig. S12 - Cell morphology images of BSPNO cross-linked cells at different time.**

**a**. Images of cells cross-linked with BSPNO for different time.

**b**. The cells that we counted to examine the diameter.

**c**. Average ratio of the cell diameter compared with 3 min cross-linking. Cell number was 15. Data are average ± s.e.m.

**Supplementary Fig. S13 - Statistics of the cross-linked sites of BSA identified from BSPNO cross-linking at different time.**

**Supplementary Fig. S14 - Mapping the cross-links identified by 5 min BSPNO cross-linking but not identified by 60 min BSPNO cross-linking on the BSA structure.**

There were five commonly identified cross-linked sites within the scope of 27 ± 6 Å Cα–Cα distances from two repeated experiments identified by 5 min BSPNO cross-linking, but not identified by 60 min BSPNO cross-linking. These sites were mapped to the BSA structure (PDB code: 3V03).

**Supplementary Fig. S15 – Exploration of cross-linked peptides enrichment with standard peptide.**

The effect of enrichment was optimized by adding a serial of the molar weight series of the click-chemistry reagent azide-diazo-biotin (ADB) to the cross-linked peptide (Ac-IEAEKGR) at two-, four- and eight-fold times the molar number of NHS groups of BSPNO and allowing reaction for 2 h at room temperature. Then, the obtained modified peptides were analyzed with a MALDI-TOF/TOF mass spectrometer.

**A.** MALDI-TOF spectra of Ac-IEAEKGR (I); cross-linked with BSPNO (II); and clicked with ADB (III). **B.** Spectra of ADB-modified cross-linked peptides with molar ratios of 1:2, 1:4, and 1:8. **C.** The clicked peptides (I) were enriched with streptavidin beads. After enrichment, the reaction supernatant had no residual modified peptides (II); then, the cross-linked peptides were released with $Na_2S_2O_4$ from streptavidin beads and subjected to MS analysis (III).

**Supplementary Fig. S16 - SDS-PAGE for evaluating BSPNO cross-linking efficiency with BSA.**

BSA (1 mg/mL) in 20 mM HEPES containing 50 mM NaCl and 1.5 mM $MgCl_2$, pH 7.4, was mixed with different concentrations of BSPNO at mass ratios of 16:1, 8:1, 4:1, 2:1, and 1:1, and then the cross-linking reaction was quenched by 2 M ABC. Two micrograms of cross-linked protein was loaded onto SDS-PAGE gels to examine the cross-linking efficiency. A 2:1 ratio was used for *E. coli* lysate proteins cross-linking.

**Supplementary Fig. S17 - MS2 spectra of DSS and BSPNO cross-linked peptides.** Top: DSS; bottom: BSPNO. The identified fragment ions are reported on the sequences.

**Supplementary Fig. S18 - Percentage distribution of the number of MS2 spectra matched to BSPNO and DSS cross-linked peptides (PSMs).**

**Supplementary Fig. S19 - Comparison of the protein copy number distribution between our XL proteome and the MS proteome of human cells.** Protein copy number was determined by shotgun proteomics in Jensen et al. (2).

**Supplementary Fig. S20 - Gene Ontology (GO) analysis of the interacting proteins.**

**Supplementary Fig. S21 - Typical subnetworks profiling of living cell cross-linking.** (A) Histone proteins interaction network. The proteins identified with direct interactions with histone proteins were illustrated as individual nodes. The histone proteins were grouped by H1, H2A, H2B, H3 and H4 shown as round rectangle. The interacting proteins were color coded based on the function annotation. (B) HSPs interaction network. The proteins identified with direct interactions with HSPs were illustrated as individual nodes. The HSP proteins were grouped by Hsp10, Hsp60, Hsp70 and Hsp90 shown as round rectangle. The interacting proteins were color coded based on the function annotation. (C) TFs interaction network. The proteins identified with direct interactions with TFs were illustrated as individual nodes. The TFs were colored green as diamond and the interacting proteins were colored grey as ellipse.

**Supplementary Fig. S22 - In vitro cross-linking subnetworks.**

(A) Histone proteins interaction network. (B) HSPs interaction network. (C) TFs interaction network.

**Supplementary Fig. S23 - Intensity distribution of the PPIs among protein copy number with reference to the distribution of the MS proteome of human cells.** Protein copy number was determined by shotgun proteomics in the data of Jensen et al. (2).

**Supplementary Fig. S24 - Overlap of our identified PPIs with the integrated PPIs databases of STRING and BioGRID and the selected published human CXMS studies.** The selected human CXMS studies were Wheat et al. 2021 (3), Chavez et al. 2019 (4), Liu et al, 2017 (5).

**Supplementary Fig. S25 - Mapping the cross-links to structural architecture.** a. Human mitochondrial chaperonin symmetrical "football" complex (PDB: 4PJ1), the single interprotein cross-link is shown as green lines within 27 Å. b. Human mitochondrial acetoacetyl-CoA thiolase (PDB: 2F2S). c. Actin (PDB: 3BYH). The same comments as on Fig. 3D apply.

**Supplementary Fig. S26 - Immunofluorescence and Western blot analysis of PTEN and the interaction proteins.**

**Supplementary Fig. S27 - Ingenuity canonical pathway analysis of cytoplasm PTEN interaction proteins.** (The red columns are all closely related to PI3K-AKT signaling pathway.)

**Supplementary Fig. S28 - Ingenuity canonical pathway analysis of nucleus PTEN interaction proteins.** (The red columns are all closely related to DNA damage and repair pathway.)

**Supplementary Fig. S29 - Ingenuity canonical pathway analysis of PTEN nuclear translocation interaction proteins.** (The red columns are all closely related to cell apoptosis.)

**Supplementary Fig. S30 - Types and sequences of PTEN isoforms.**

**Supplementary Fig. S1**

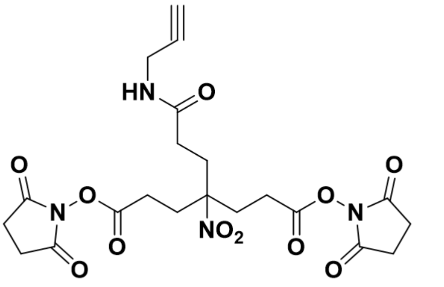


**BSPNO, arm length: 8.9 Å**

**Supplementary Fig. S2**

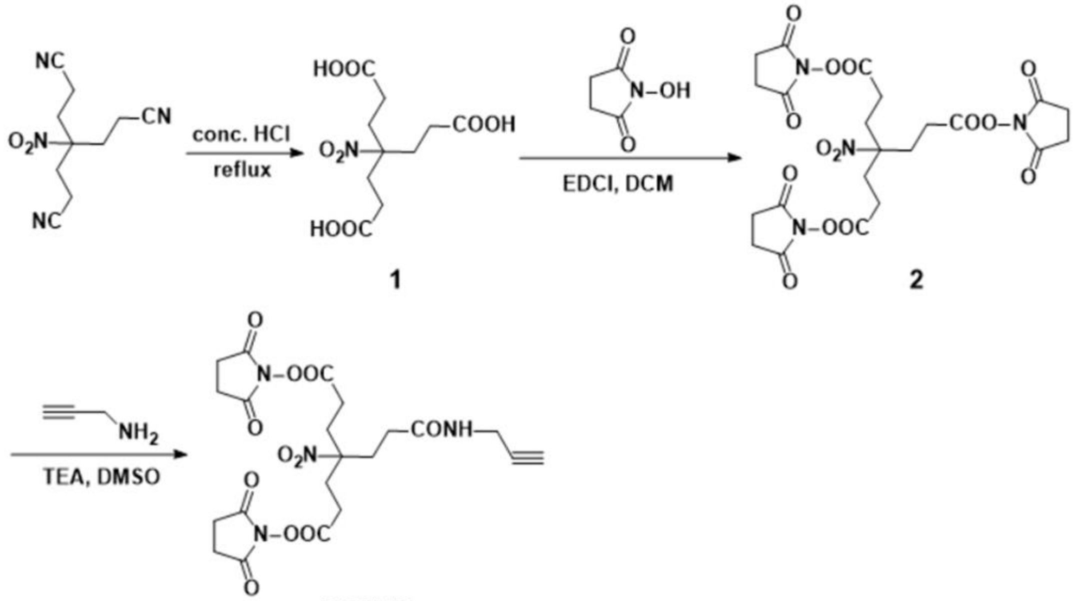


BSPNO

**Supplementary Fig. S3**

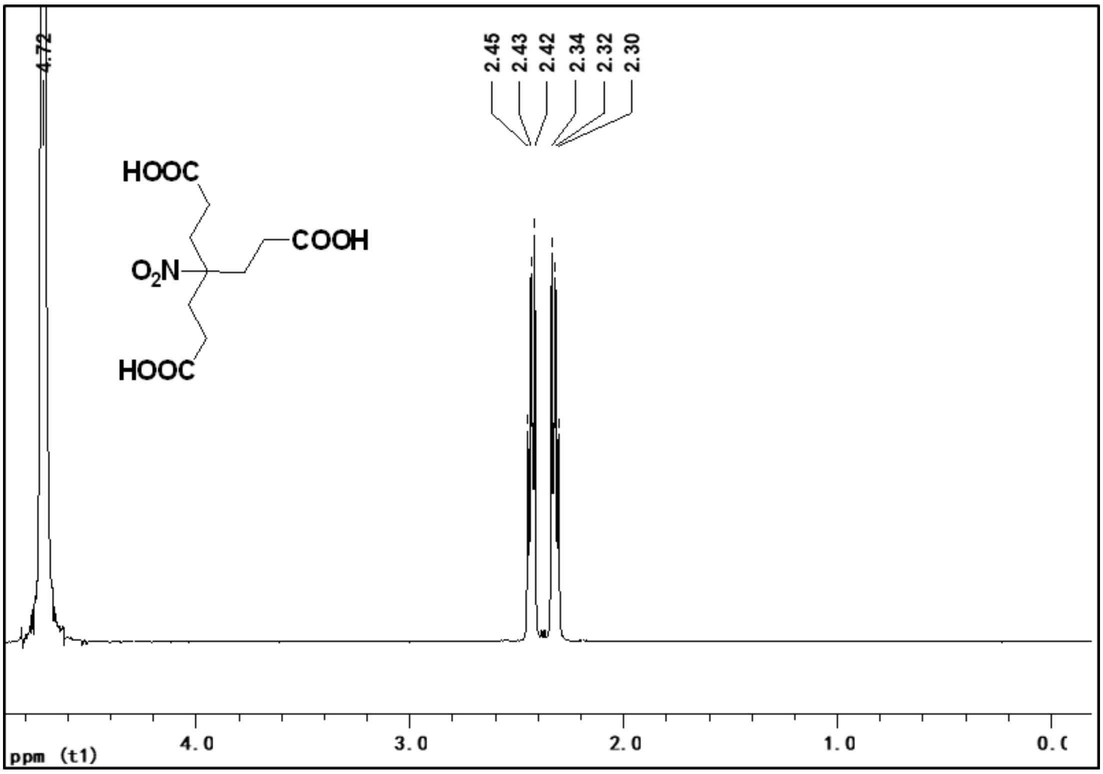

**Supplementary Fig. S4**

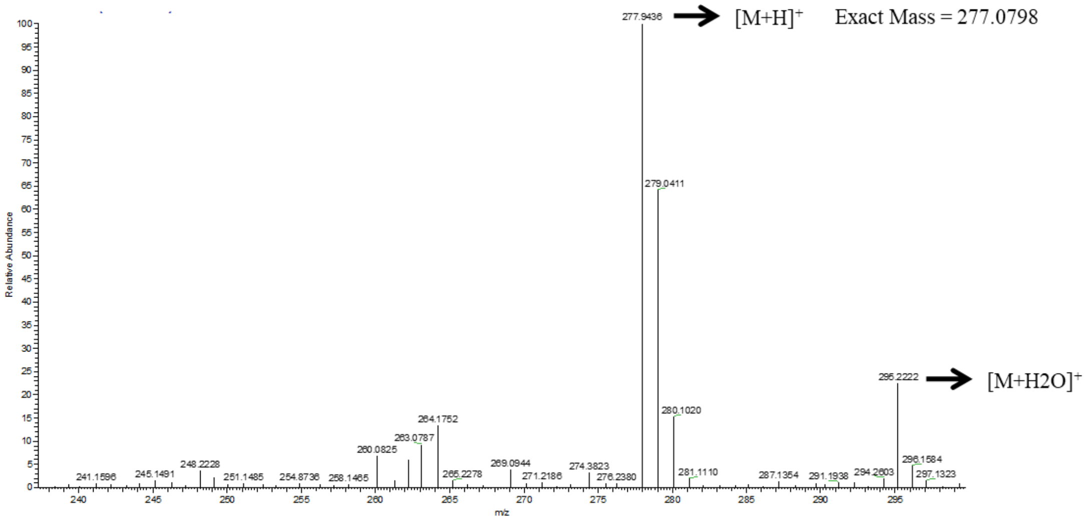

**Supplementary Fig. S5**

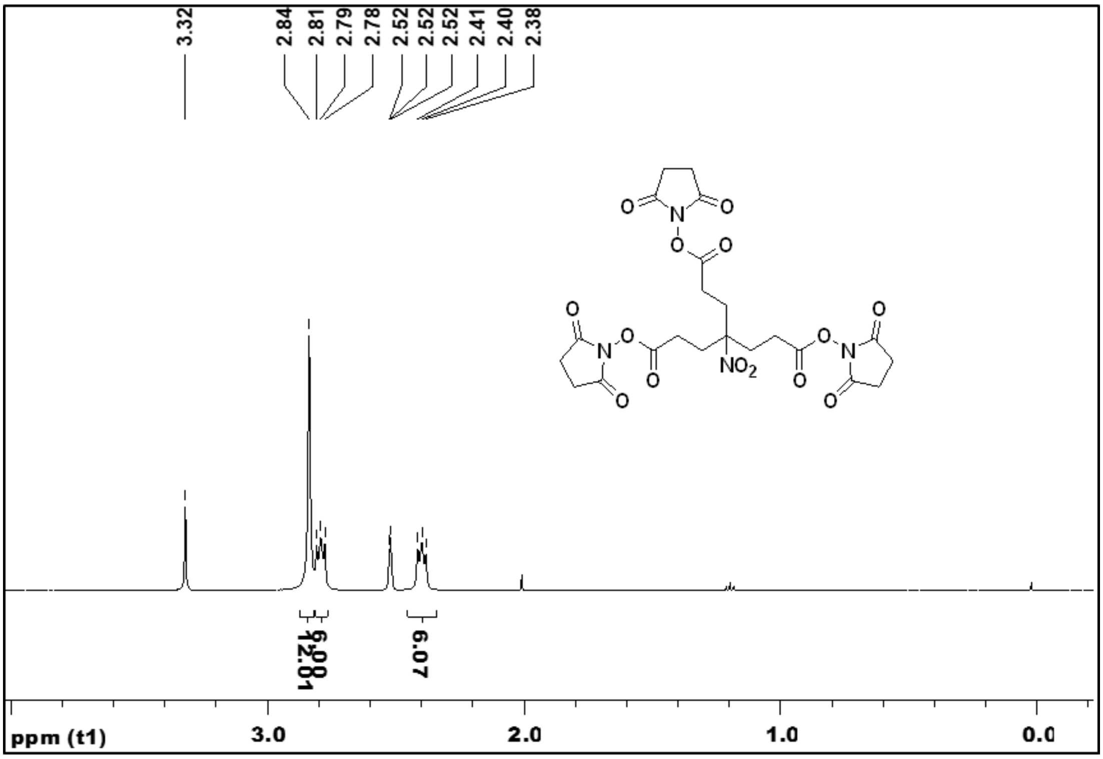

**Supplementary Fig. S6**

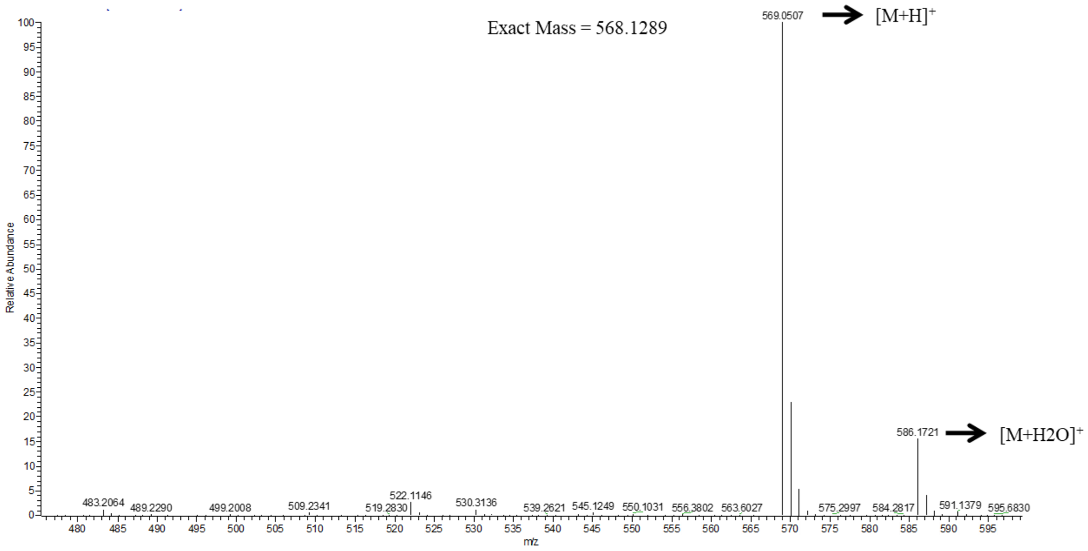

**Supplementary Fig. S7**

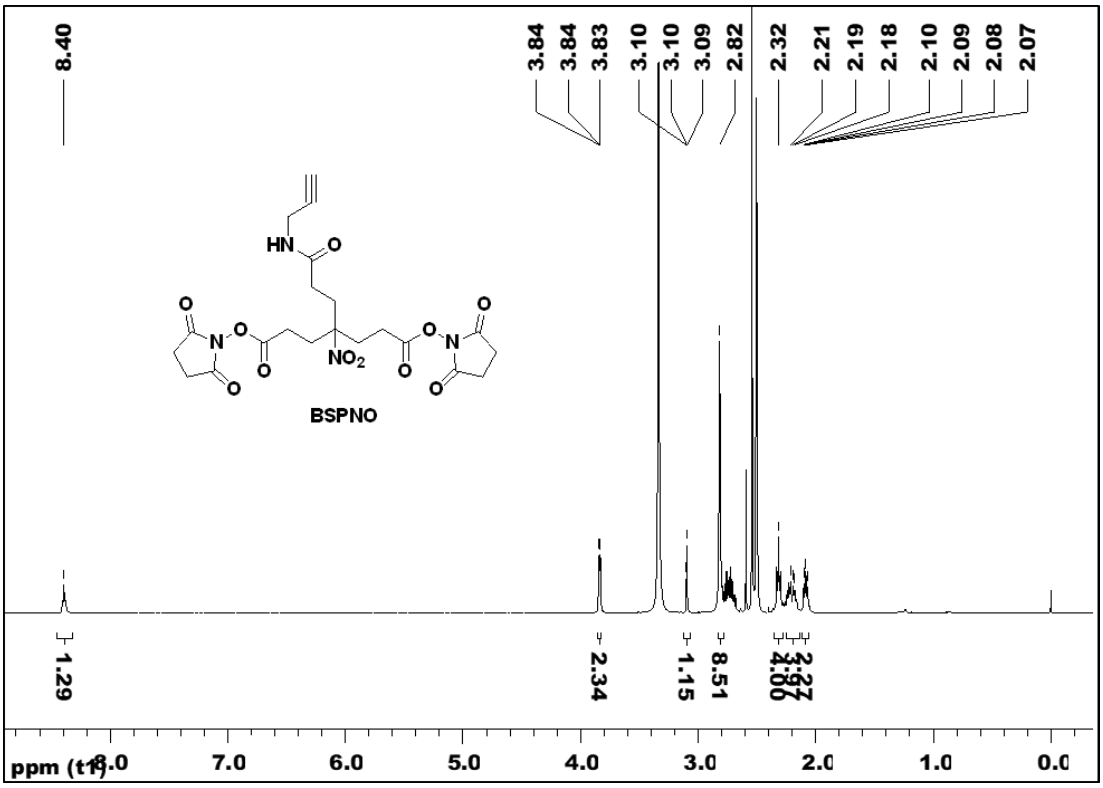

**Supplementary Fig. S8**

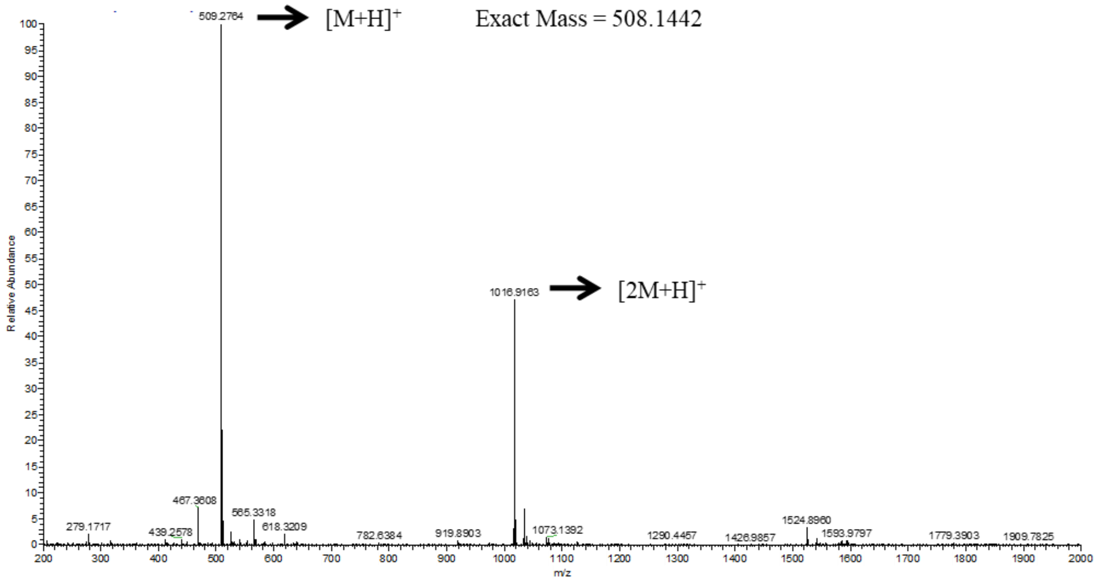

**Supplementary Fig. S9**

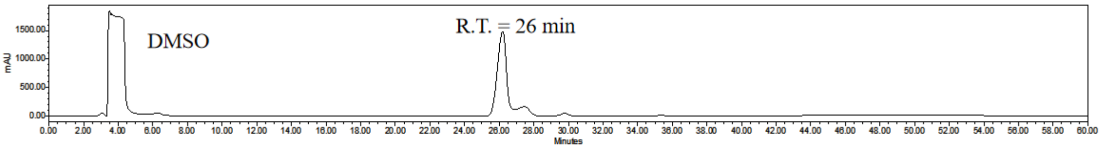

**Supplementary Fig. S10**

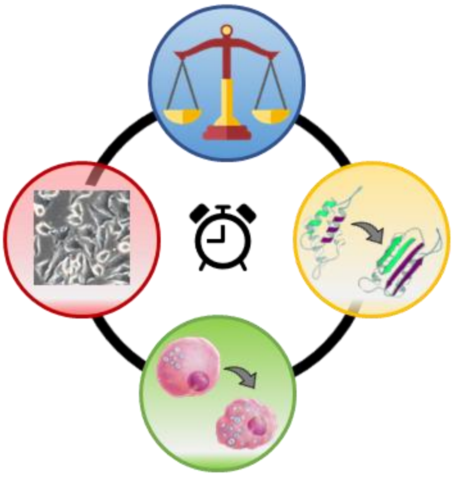

**Supplementary Fig. S11**

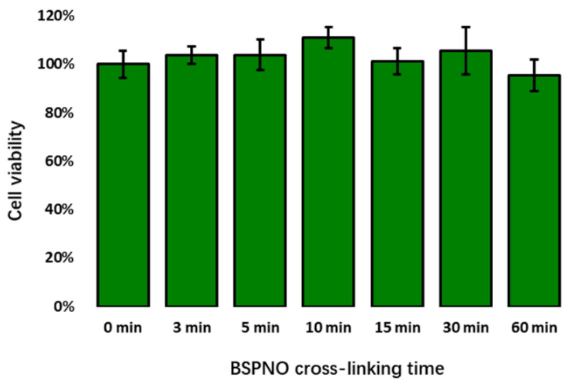

**Supplementary Fig. S12**

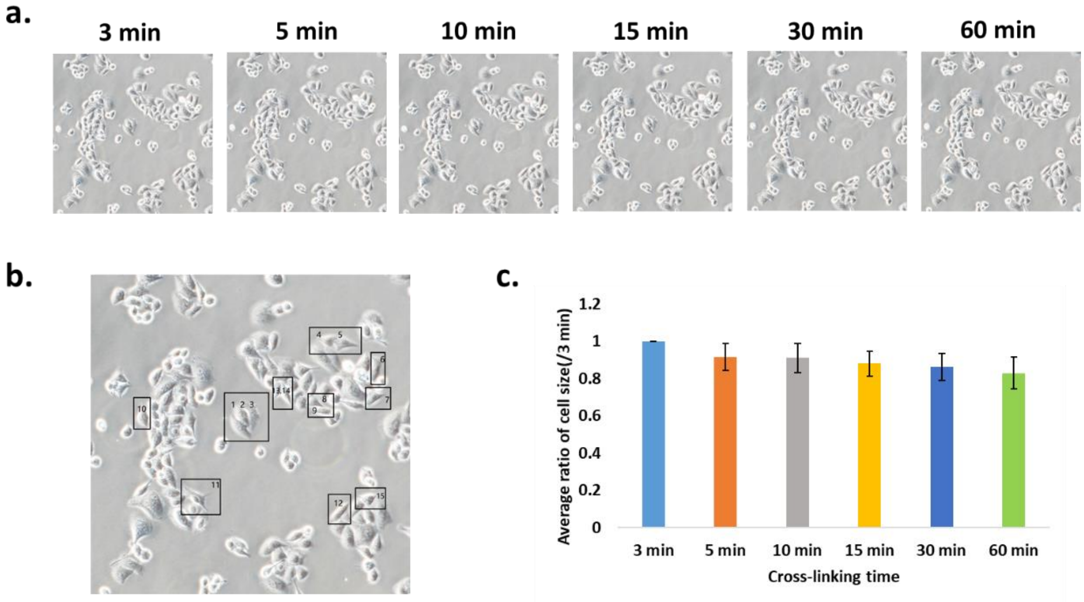

#### Supplementary Fig. S13

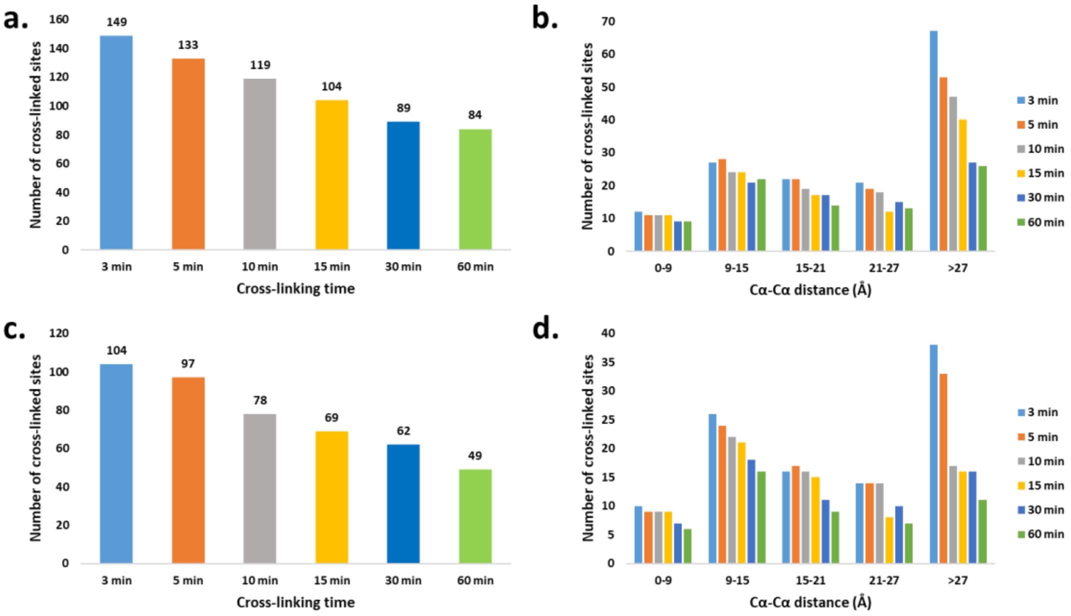

**Supplementary Fig. S14**

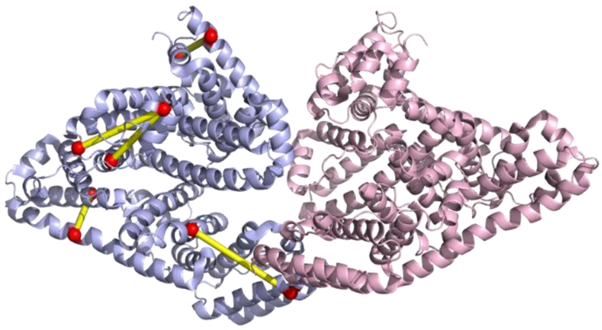

**Supplementary Fig. S15**

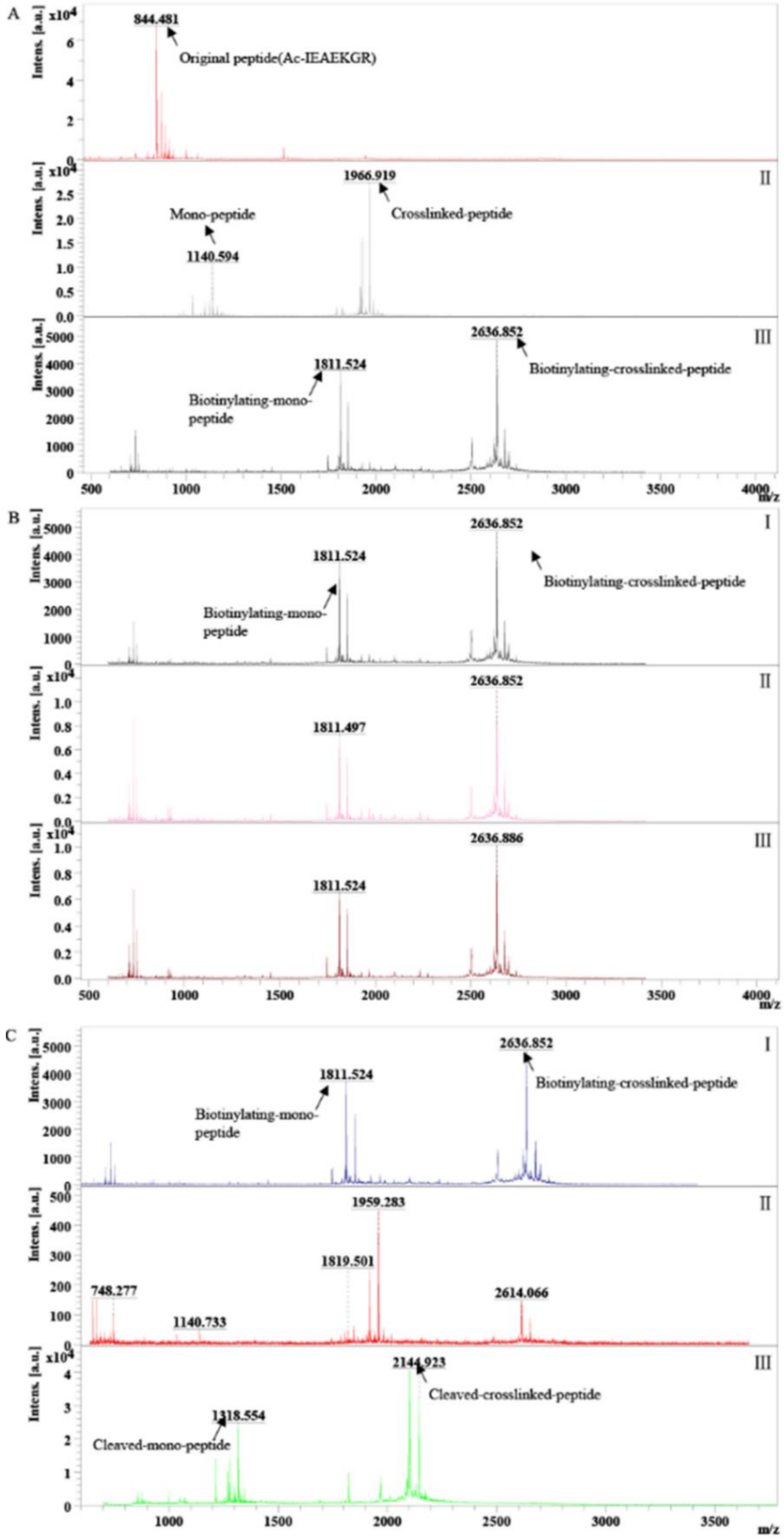

**Supplementary Fig. S16**

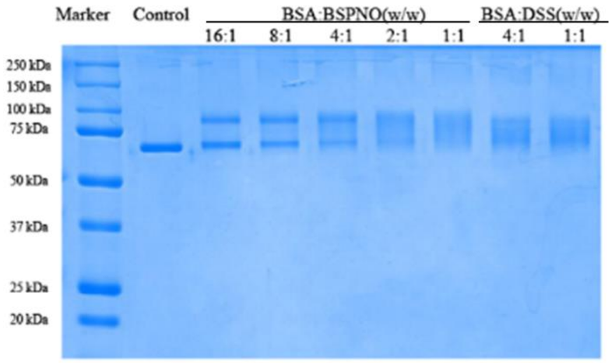

#### Supplementary Fig. S17

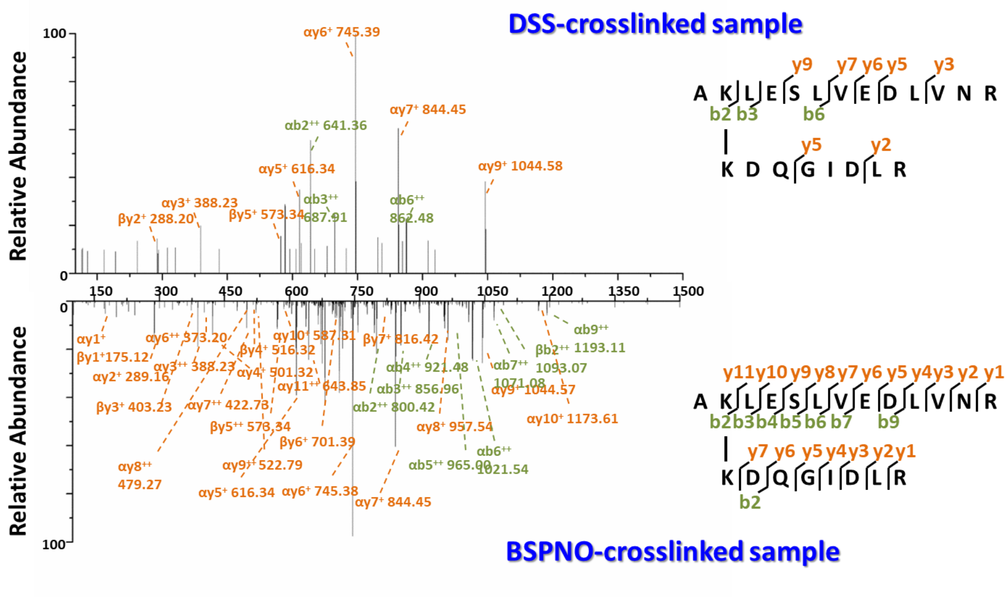

#### Supplementary Fig. S18

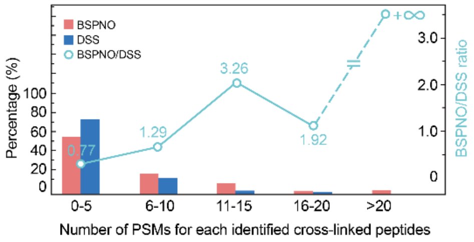

**Supplementary Fig. S19**

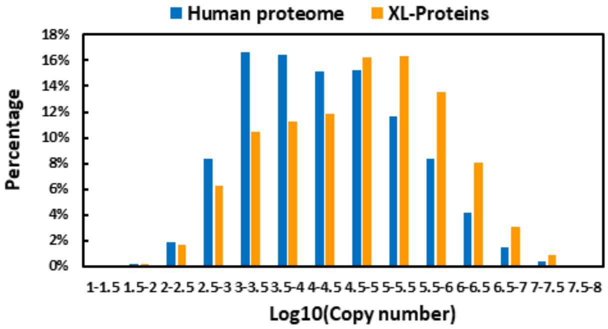

#### Supplementary Fig. S20

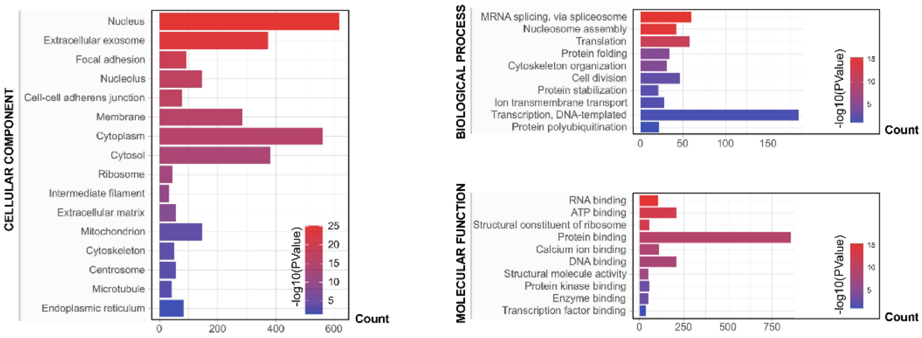

**Supplementary Fig. S21**

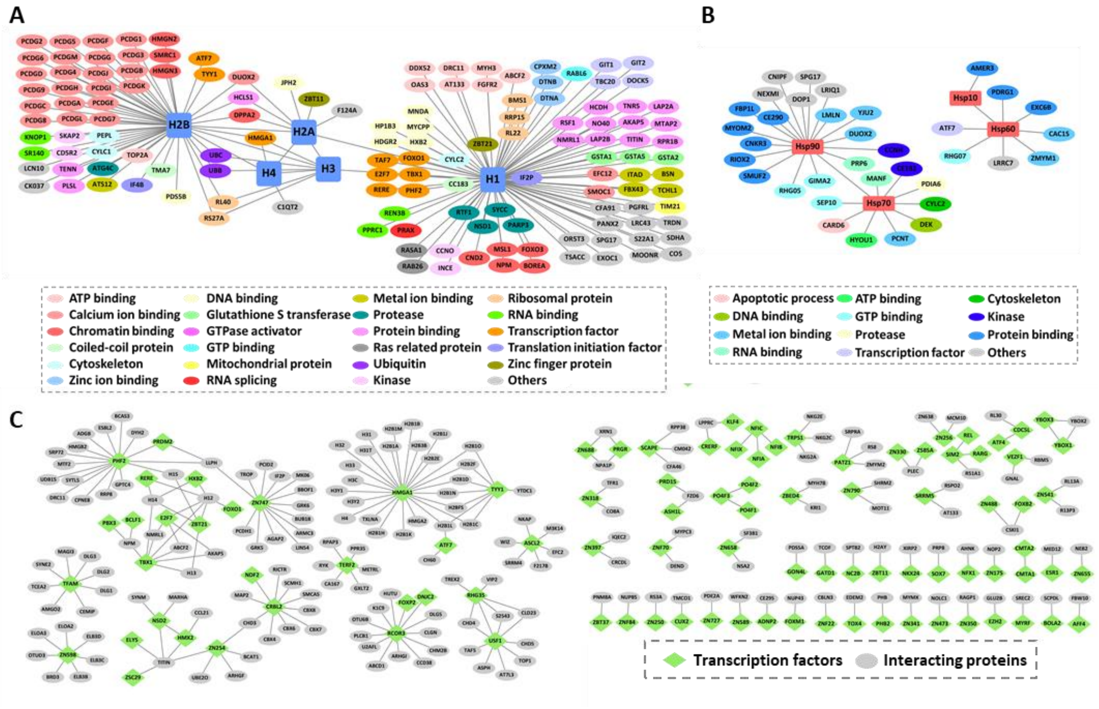

**Supplementary Fig. S22**

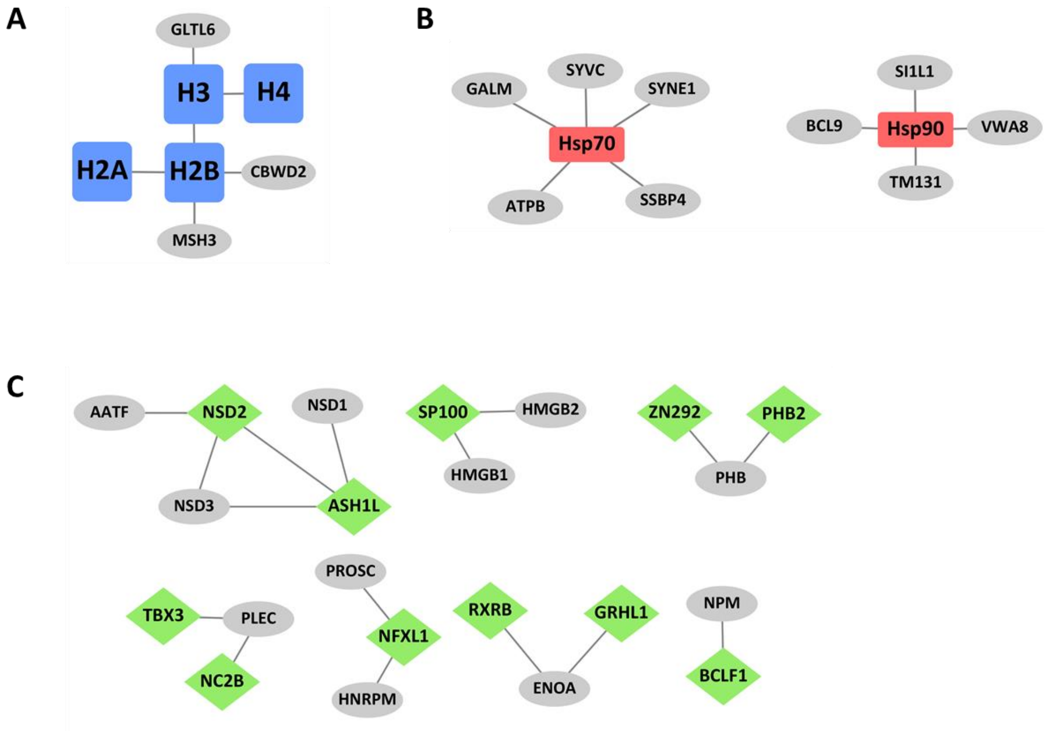

**Supplementary Fig. S23**

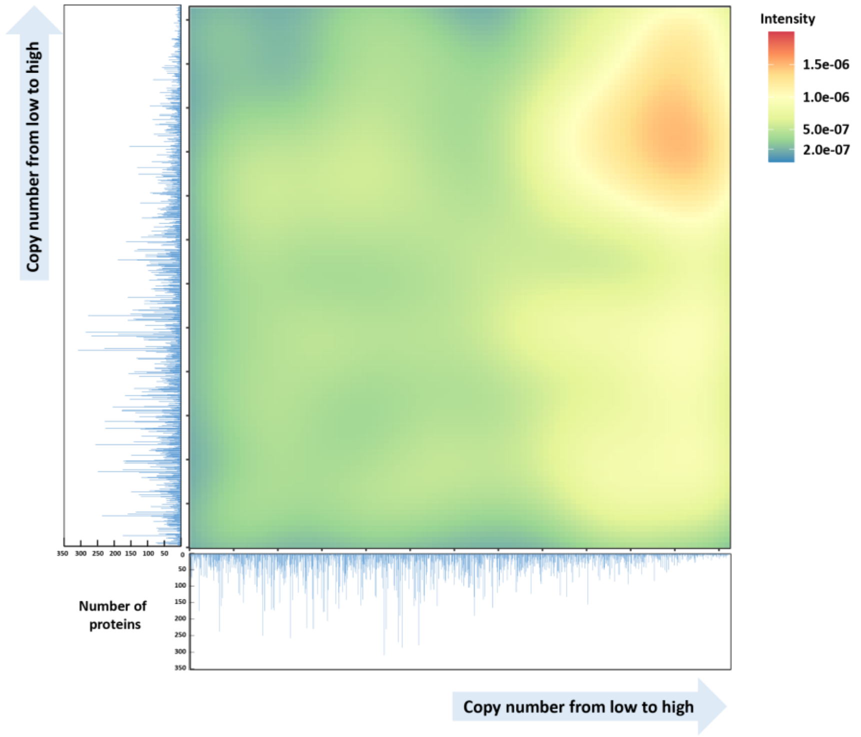

**Supplementary Fig. S24**

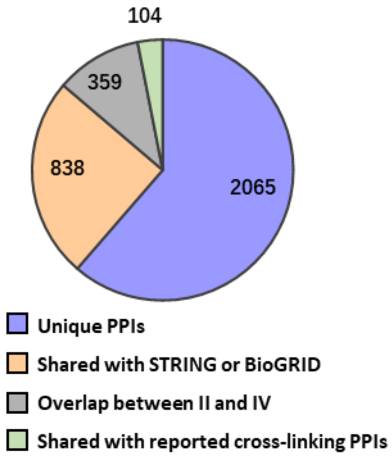

**Supplementary Fig. S25**

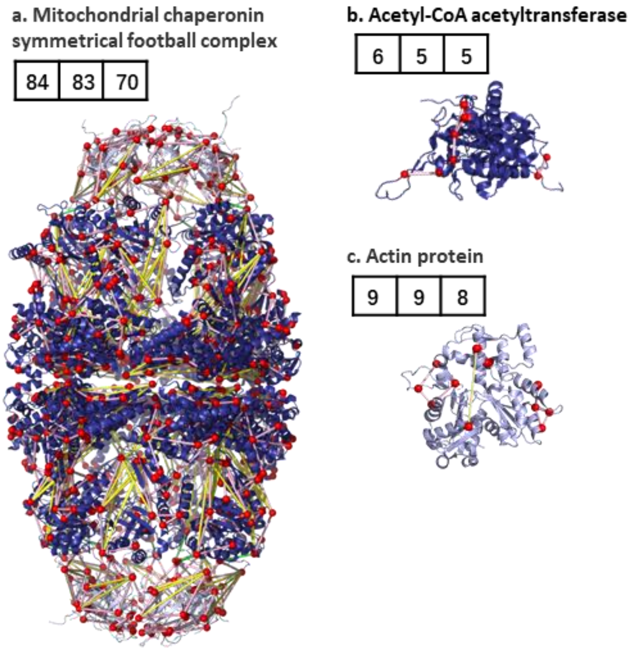

Supplementary Fig. S26

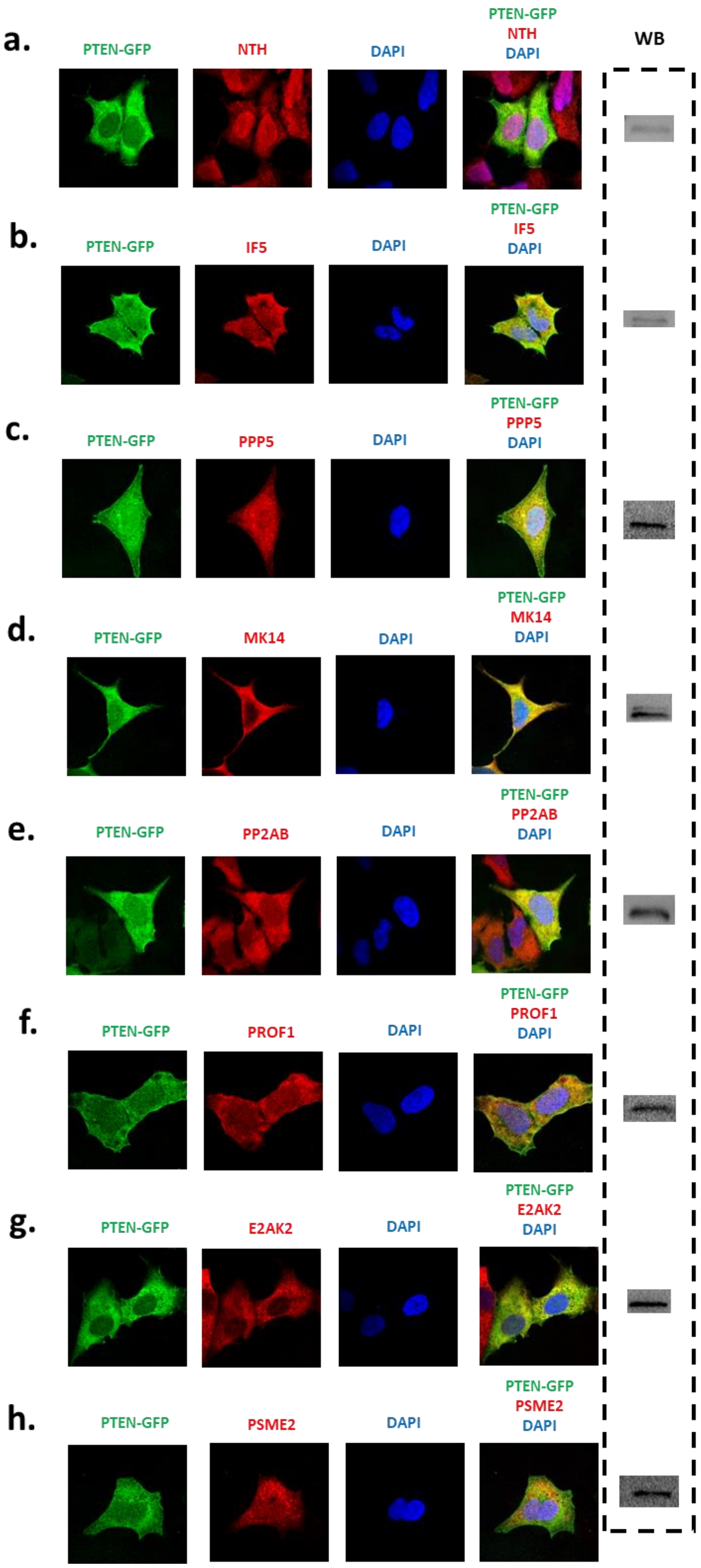

**Supplementary Fig. S27**

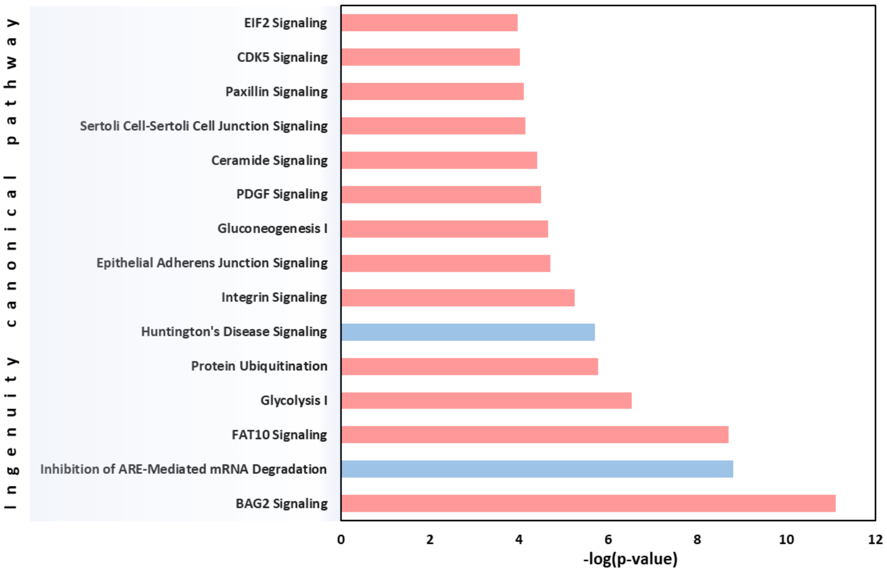

**Supplementary Fig. S28**

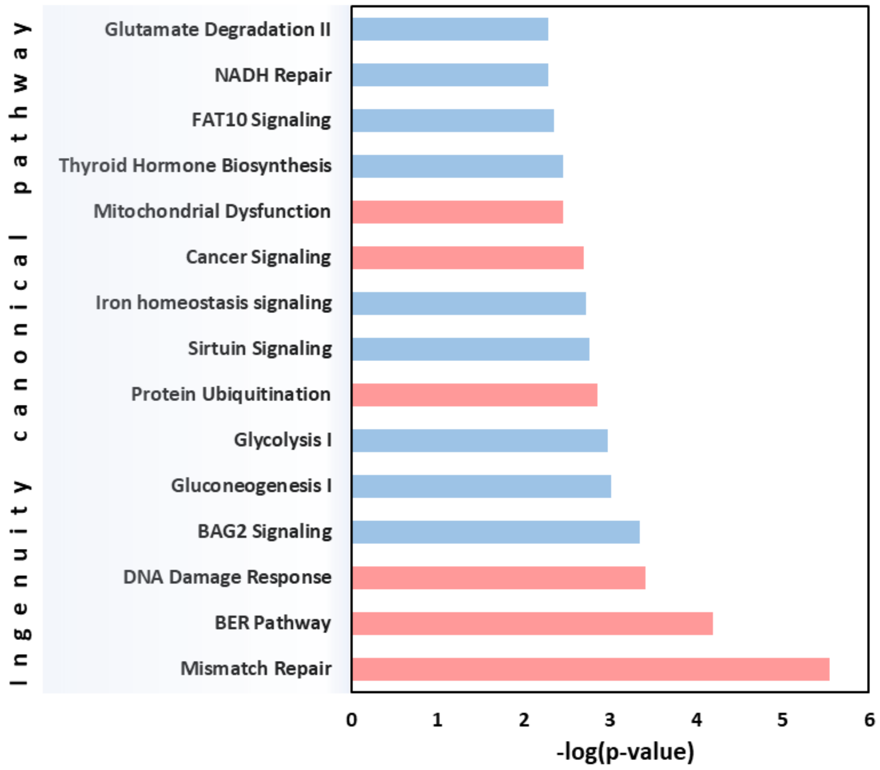

**Supplementary Fig. S29**

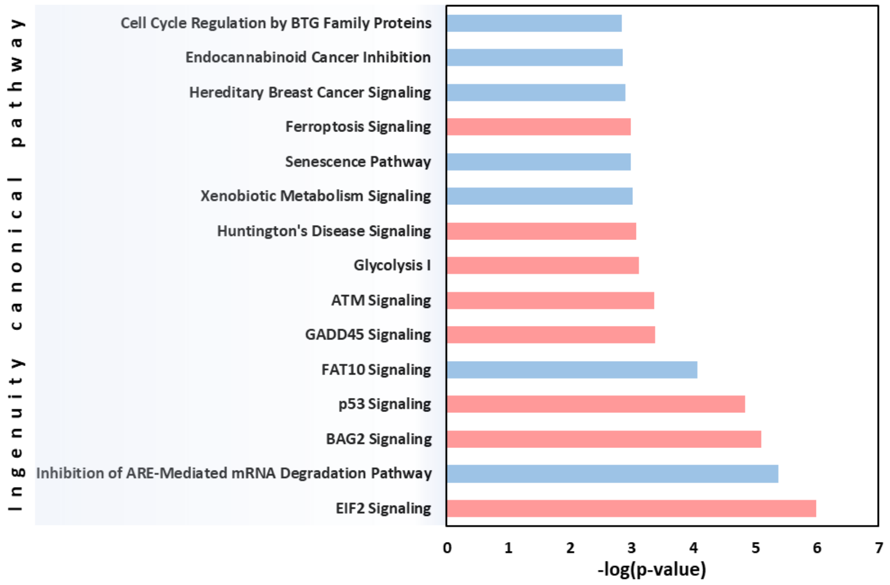

**Supplementary Fig. S30**

**P60484-1**, PTEN, **isoform1**, the canonical sequence
MTAIIKEIVSRNKRRYQEDGFDLDLTYIYPNIIAMGFPAERLEGVYRNNIDDVVRFLDSK
HKNHYKIYNLCAERHYDTAKFNCRVAQYPFEDHNPPQLELIKPFCEDLDQWLSEDDNHVA
AIHCKAGKGRTGVMICAYLLHRGKFLKAQEALDFYGEVRTRDKKGVTIPSQRRYVYYYSY
LLKNHLDYRPVALLFHKMMFETIPMFSGGTCNPQFVVCQLKVKIYSSNSGPTRREDKFMY
FEFPQPLPVCGDIKVEFFHKQNKMLKKDKMFHFWVNTFFIPGPEETSEKVENGSLCDQEI
DSICSIERADNDKEYLVLTLTKNDLDKANKDKANRYFSPNFKVKLYFTKTVEEPSNPEAS
SSTSVTPDVSDNEPDHYRYSDTTDSDPENEPFDEDQHTQITKV

**P60484-2**, PTEN-long, **isoform2**, produced by alternative initiation at a CTG start codon of isoform 1.
The sequence of this isoform differs from the canonical sequence as follows:
1-1: M → MERGGEAAAA...FFFSHRLPDM
**MERGGEAAAAAAAAAAAPGRGSESPVTISRAGNAGELVSPLLLPPTRRRRRRHIQGPGPV**
**LNLPSAAAAPPVARAPEAAGGGSRSEDYSSSPHSAAAAARPLAAEEKQAQSLQPSSSRRS**
**SHYPAAVQSQAAAERGASATAKSRAISILQKKPRHQQLLPSLSSFFFSHRLPD**MTAIIKE
IVSRNKRRYQEDGFDLDLTYIYPNIIAMGFPAERLEGVYRNNIDDVVRFLDSKHKNHYKI
YNLCAERHYDTAKFNCRVAQYPFEDHNPPQLELIKPFCEDLDQWLSEDDNHVAAIHCKAG
KGRTGVMICAYLLHRGKFLKAQEALDFYGEVRTRDKKGVTIPSQRRYVYYYSYLLKNHLD
YRPVALLFHKMMFETIPMFSGGTCNPQFVVCQLKVKIYSSNSGPTRREDKFMYFEFPQPL
PVCGDIKVEFFHKQNKMLKKDKMFHFWVNTFFIPGPEETSEKVENGSLCDQEIDSICSIE
RADNDKEYLVLTLTKNDLDKANKDKANRYFSPNFKVKLYFTKTVEEPSNPEASSSTSVTP
DVSDNEPDHYRYSDTTDSDPENEPFDEDQHTQITKV

**P60484-3**, **isoform3**
The sequence of this isoform differs from the canonical sequence as follows:
55-70: RFLDSKHKNHYKIYNL → S
165-190: GVTIPSQRRYVYYYSYLLKNHLDYRP → ADPTGGIPDKGIIVIGDGSSMDVIAP
191-403: Missing.
MTAIIKEIVSRNKRRYQEDGFDLDLTYIYPNIIAMGFPAERLEGVYRNNIDDVV**S**CAERH
YDTAKFNCRVAQYPFEDHNPPQLELIKPFCEDLDQWLSEDDNHVAAIHCKAGKGRTGVMI
CAYLLHRGKFLKAQEALDFYGEVRTRDKK**ADPTGGIPDKGIIVIGDGSSMDVIAP**
